# Loss of a subunit of vacuolar ATPase identifies unexpected biological signatures of reduced organelle acidification *in vivo*

**DOI:** 10.1101/2020.11.30.397463

**Authors:** Pottie Lore, Van Gool Wouter, Vanhooydonck Michiel, Hanisch Franz-Georg, Goeminne Geert, Rajkovic Andreja, Coucke Paul, Sips Patrick, Callewaert Bert

## Abstract

The inability to maintain a strictly regulated endo(lyso)somal acidic pH through the proton-pumping action of the vacuolar-ATPases has been associated with various human diseases including heritable connective tissue disorders, neurodegenerative diseases and cancer. Multiple studies have investigated the pleiotropic effects of reduced acidification *in vitro*, but the mechanisms elicited by impaired endo(lyso)somal acidification *in vivo* remain poorly understood. Here, we show that loss of *atp6v1e1b* in zebrafish leads to early mortality, associated with craniofacial dysmorphisms, vascular anomalies, cardiac dysfunction, hypotonia and epidermal structural defects, reminiscent of the phenotypic manifestations in cutis laxa patients carrying a defect in the *ATP6V1E1* gene. Mechanistically, we found that *in vivo* genetic depletion of *atp6v1e1b* leads to N-glycosylation defects and reduced maturation of endosomal and lysosomal vesicles, but retains the hypoxia-mediated response. In order to gain further insights into the processes affected by aberrant organelle acidification, we performed an untargeted analysis of the transcriptome and metabolome in early *atp6v1e1b*-deficient larvae. We report multiple affected pathways including but not limited to oxidative phosphorylation, sphingolipid and fatty acid metabolism with profound defects in mitochondrial respiration. Taken together, our results identify new complex biological effects of reduced organelle acidification *in vivo*, which likely contribute to the multisystemic manifestations observed in disorders caused by v-ATPase deficiency.

## Introduction

The intracellular pH of specific subcellular compartments is maintained by the proton-pumping action of the vacuolar-ATPases (v-ATPase), thereby regulating a range of molecular processes including activation of enzyme activity, protein folding, vesicle trafficking, and support of organelle function and integrity. V-ATPases consist of ubiquitously expressed multisubunit complexes, composed of a membrane-associated V_0_ domain and a cytosolic (catalytic) V_1_ domain. Hydrolysis of ATP by the complex generates proton pumping activity against a concentration gradient in the lumen of endo(lyso)somes, secretory vesicles, Golgi apparatus and across the plasma membrane of specialized cells including osteoclasts, epididymal clear cells and renal epithelial intercalating cells [1].

Various human diseases are linked to defects in genes encoding subunits of the v-ATPase, including neurological disorders (e.g. encephalopathy with epilepsy (*ATP6V1A* [MIM: 618012]), X-linked mental retardation Hedera type (*ATP6AP2* [MIM: 300423]), and X-linked Parkinson Disease with spasticity (*ATP6AP2* [MIM: 300911])), congenital disorders of glycosylation (*ATP6AP1* [MIM: 300972] and *ATP6AP2* [MIM: 301045]), disorders in which the secretion of protons in the extracellular environment is disrupted (e.g. distal renal tubular acidosis and hearing loss (*ATP6V1B1* [MIM: 192132] and *ATP6V0A4* [MIM: 605239]), syndromic osteopetrosis (*ATP6V0A3* [MIM:259700]), Zimmermann-Laband syndrome (*ATP6V1B2* [MIM: 616455]), dominant deafness-onychodystrophy syndrome (*ATP6V1B2* [MIM: 124480])) and connective tissue disorders (e.g. autosomal recessive (AR) cutis laxa (CL) syndrome type 2A (ATP6V0A2 [MIM: 219200]), ARCL type 2C (ATP6V1E1 [MIM: 617402]), and ARCL type 2D (ATP6V1A [MIM: 617403])) [2–12]. ARCL type 2 shows variable glycosylation abnormalities [10, 13, 14], and is characterized by skin wrinkles or loose redundant skin folds, variable level of mental impairment, delayed neuromotor development, sensorineural hearing loss, skeletal abnormalities, facial dysmorphology, hypotonia, and cardiopulmonary involvement including pneumothorax, hypertrophic cardiomyopathy, and aortic root dilation [10, 11, 15, 16]. Finally, v-ATPases have been implicated in cancer growth and metastasis in a variety of contexts [17]. Higher v-ATPase expression and relocation to the plasma membrane is observed in proliferating cancer cells including breast, prostate, liver, pancreatic, melanoma, and esophageal cancers [18]. V-ATPase activity in the plasma membrane promotes cell survival since the resulting alkalization of the intracellular environment supports the Warburg effect by reducing the excess acid load caused by glycolytic lactic acid production [19].

Intracellular v-ATPases further regulate key physiological cell signaling pathways. V-ATPase are involved in downstream Notch signaling [20–23], Wnt signaling [18, 24–29], activation of mechanistic target of rapamycin complex 1 (mTORC1), and AMP-activated protein kinase (AMPK) signaling [17]. Recent studies have highlighted a novel mechanism linking v-ATPase inhibition, hypoxia-inducible factor (HIF) signaling and iron deficiency. Using *in vitro* experiments, it was shown that genetic and pharmacological inhibition of v-ATPases impairs endo(lyso)somal acidification which leads to defective cellular iron uptake via the transferrin system. Iron depletion results in reduced prolyl hydroxylase activity, leading to stabilization of HIF and activation of HIF-dependent signaling mechanisms [30–32]. This raises the question whether *in vivo* v-ATPase deficiency impairs iron homeostasis and perturbates HIF-mediated response, and if this represents the major mode of action in genetic disorders related to defects in v-ATPase subunits.

Complete loss of the murine proteolipid 16 kDa subunit of the v-ATPase V_0_ complex is embryonically lethal [33]. Since external fertilization and transparent development allow us to study embryonic development in zebrafish, we modelled genetically depleted *atp6v1e1b* zebrafish. Despite early mortality, this model mimics ARCL type 2C with hypotonia, craniofacial abnormalities, vascular anomalies, and an altered dermal connective tissue structure. Untargeted analysis of the transcriptome, metabolome, and lipidome furthermore indicates alterations in oxidative phosphorylation, fatty acid metabolism, and sphingolipid metabolism in this *in vivo* model of v-ATPase deficiency.

## Results

### Genetic loss of *atp6v1e1b* in zebrafish leads to early mortality and mimics the human phenotype of *ATP6V1E1*-related cutis laxa syndrome

We genetically depleted *atp6v1e1b,* a subunit of the v-ATPase that connects the transmembrane domain with the cytosolic domain, in zebrafish to identify pleiotropic manifestations of reduced organelle acidification *in vivo.* We generated a zebrafish line harboring a two base-pair insertion followed by a three base-pair deletion in exon 5 of *atp6v1e1b* (*atp6v1e1b*^cmg78/+^), c.334insGG; c.337-340delCGG, which led to nonsense-mediated decay (NMD) due to a premature stop codon (**Supplemental Figure 1A-B**). We further obtained a zebrafish line with viral insertion in the 5’UTR of *atp6v1e1b* (*atp6v1e1b*^hi577aTg/+^) which also led to NMD (**Supplemental Figure 1A-B**) [34, 35]. We recapitulated a pigment dilution phenotype which can be used as an early read-out. *Atp6v1e1b*-deficient zebrafish were unable to hatch and died at 3 – 5 days post fertilization (dpf) (**Supplemental Figure 1D**). Upon manual removal of the chorion, the mutant zebrafish survived until 8 dpf (**Supplemental Figure 1C**) albeit with a severe systemic phenotype. This prompted us to further investigate the phenotype of the surviving larvae to assess whether dominant features in the clinical presentation of ARCL type 2C are present in *atp6v1e1b*-deficient zebrafish. To evaluate hypotonia as a possible cause of hatching problems, we quantitatively assessed the touch-evoked escape response at 3 dpf, which was impaired in both *atp6v1e1b* mutant zebrafish lines (**Supplemental Figure 1E**). Alcian blue staining showed maxillary and mandibular hypoplasia in both mutant zebrafish which could correlate to midface hypoplasia and micrognathia in humans suffering from ARCL type 2C (**Figure 1A**). *Atp6v1e1b*-deficient zebrafish showed a misshapen and slightly retracted Meckel’s cartilage. Moreover, the length of the Meckel’s cartilage, palatoquadrate and ceratohyal structures was decreased (**Figure 1B**) and the angle between ceratohyal structures was increased (**Figure 1C**). Next, we assessed the cardiac function and vascular patterning in the *atp6v1e1b*-deficient zebrafish at the embryonic stage, using the endothelial *Tg(kdrl:eGFP)* reporter line. Vascular anomalies were present in both zebrafish lines at 5 dpf (**Figure 1E**). We discovered a structural malformation of the aortic arches, the opercular artery and the ventral aorta which were smaller and more constricted with less vascular loops in *atp6v1e1b*-deficient zebrafish. The ventral aorta segment proximal to the bulbus arteriosus was significantly dilated in *atp6v1e1b*-deficient zebrafish (**Figure 1D**). Furthermore, they showed a significantly decreased stroke volume (SV) and cardiac output (CO) at 3 dpf, indicative of cardiomyopathy (**Figure 1F**). Posterior blood flow was significantly decreased in *atp6v1e1b*^hi577aTg/hi577aTg^ but not in *atp6v1e1b*^cmg78/cmg78^ (**Figure 1F-G**). Starting from 5 dpf, mutant zebrafish developed severe pericardial edema (**Figure 1A**), indicating severe cardiovascular dysfunction.

**Figure 1:**
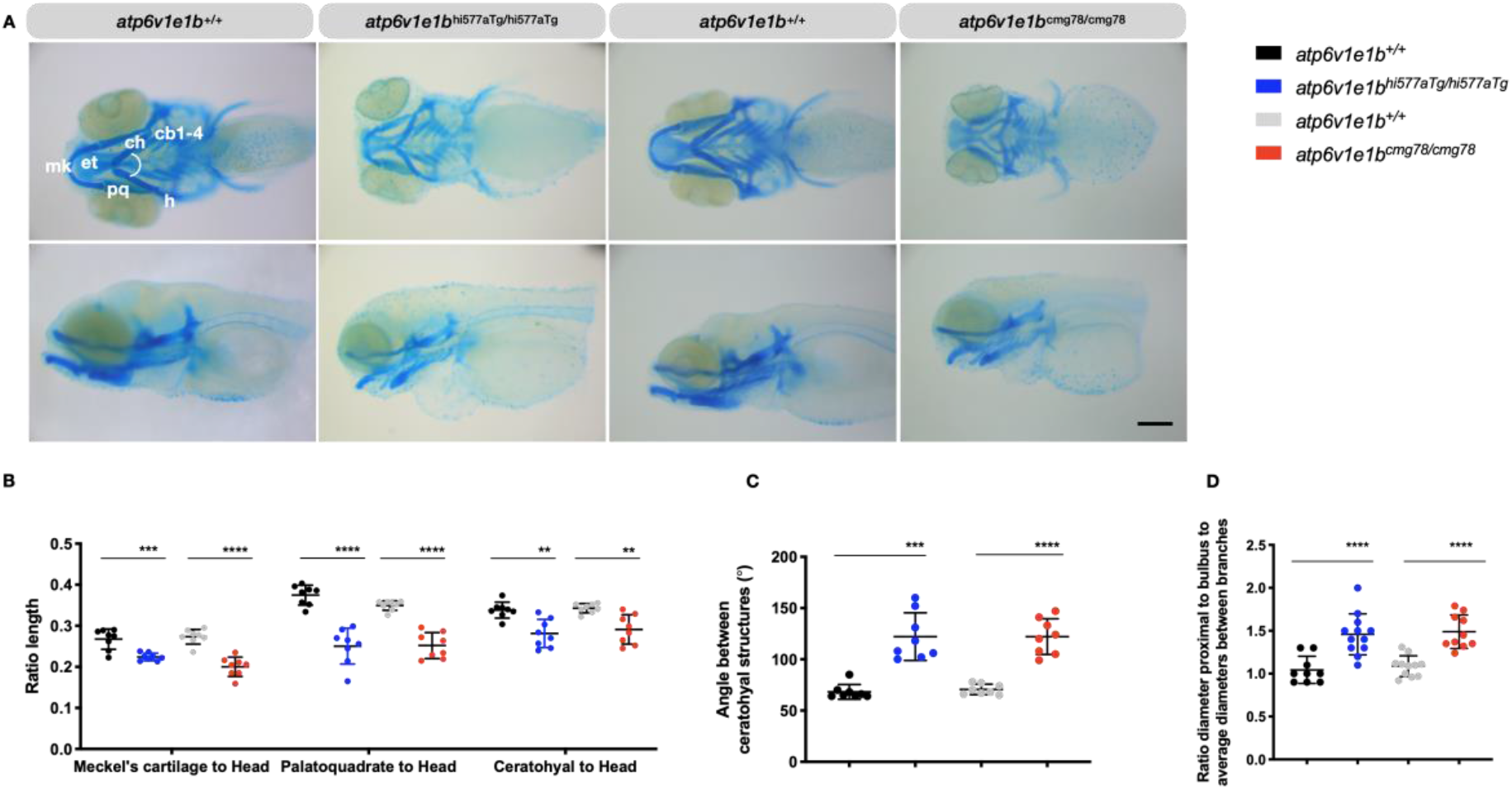
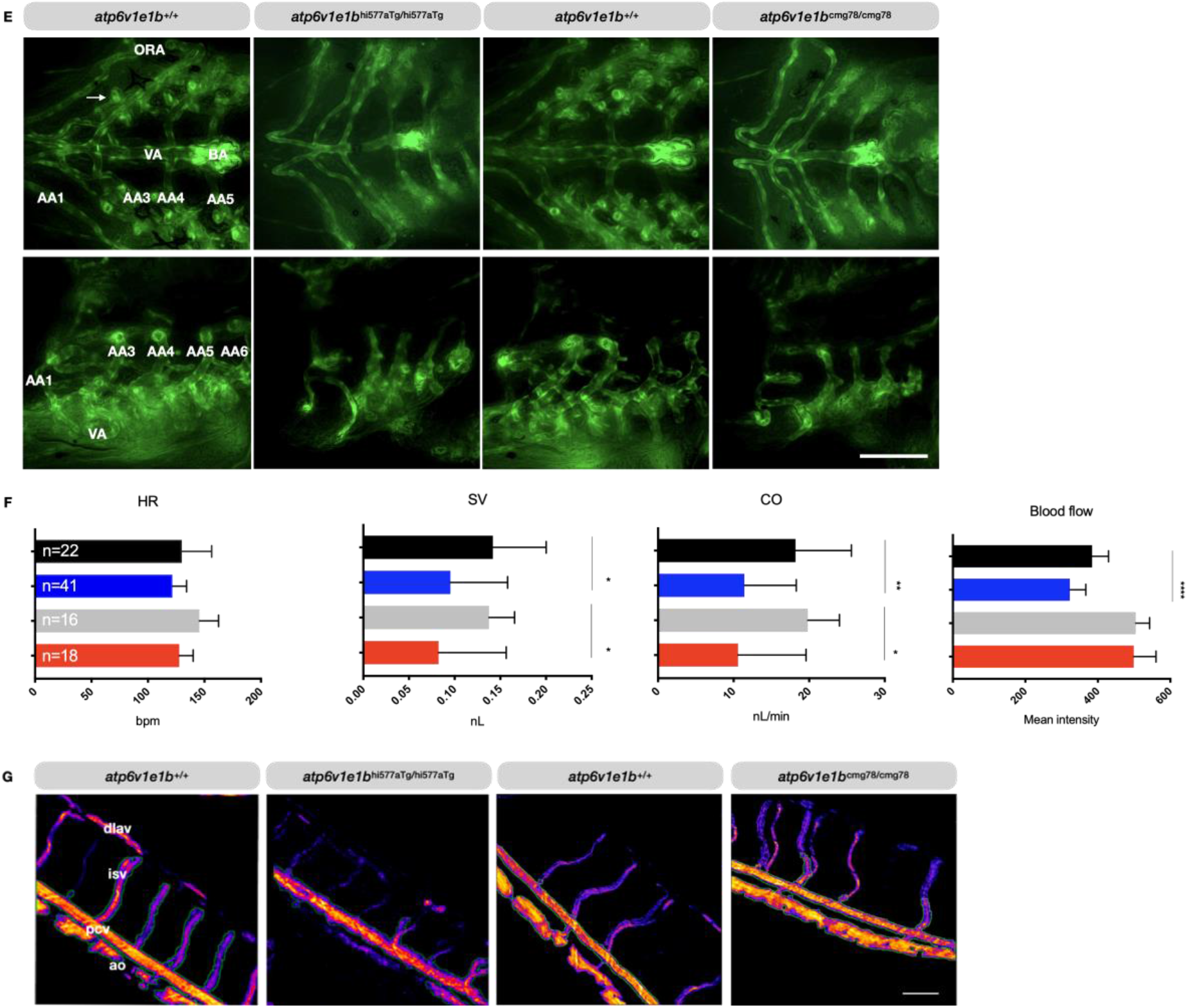
*Atp6v1e1b*-deficient zebrafish have craniofacial, vascular and cardiac abnormalities reminiscent of the human *ATP6V1E1*-related CL phenotype. Representative images are shown. (A) Ventral (top panel) and lateral views (bottom panel) of Alcian blue stained craniofacial structures at 5 dpf reveal misshapen and shorter Meckel’s cartilage (m), shorter palatoquadrate (pq) and shorter ceratohyal (ch) structure and a higher angle between ch structures in *atp6v1e1b*-deficient zebrafish. (B) The length of the individual cartilage structures was measured and normalized to the head length of the larvae. (C) Quantification of the angle between the ceratohyal structures. (D) *Atp6v1e1b*-deficient zebrafish demonstrate an increased ratio of the aortic diameter proximal to the bulbus arteriosus (BA) to the average aortic diameter between the first (AA3) and the second branchial arch (AA4) indicative for local dilatation. (E) Ventral (top panel) and lateral views (bottom panel) of endothelial cells of *atp6v1e1b*-deficient zebrafish in the *Tg(kdrl:eGFP*) background at 5 dpf indicates abnormal vascular morphology in *atp6v1e1b*-deficient zebrafish. (F) Cardiac parameter analysis based on brightfield microscopy recordings at 3 dpf. Stroke volume (SV) and cardiac output (CO) were significantly decreased in *atp6v1e1b*-deficient zebrafish, indicating cardiomyopathy. (G) Pseudo-colored processed images representing relative blood flow intensity in the trunk of 3 dpf zebrafish. Major blood vessel ROI used for quantification is marked with a green outline. cb1-4: ceratobranchial pairs 1 to 4; ch: ceratohyal; et: ethmoid plate; h: hyosymplectic; m: Meckel’s cartilage; pq, palatoquadrate; BA: bulbus arteriosus; VA: ventral aorta; AA1: mandibular arch; AA3: first branchial arch; AA4: second branchial arch; AA5: third branchial arch; AA6: fourth branchial arch; ORA: opercular artery; Ao: aorta; isv: intersegmental vessel, pcv: posterior caudal vein, dlav: dorsal longitudinal anastomotic vessel; HR: heart rhythm; SV: stroke volume; CO: cardiac output. Scale bar = 200 μm (A), scale bar = 100 μm (E and G).

Transmission electron microscopy (TEM) in skin samples at 4 dpf showed profound defects around the epidermal basement membrane structure. *Atp6v1e1b*^hi577aTg/hi577aTg^ showed zones of epidermal detachment, larger primary dermal stroma and disorganized collagenous fibrils (**Figure 2A**). Less collagen bundles were present below the basement membrane in the *atp6v1e1b*^cmg78/cmg78^. In addition, the coiled basal membrane indicates skin redundancy (**Figure 2A**). RT-qPCR analysis confirmed collagen and elastin deficiency in both mutant zebrafish larvae at 4 dpf. *Col1a1a* and *col1a2* (**Figure 2B-E**) were significantly reduced in *atp6v1e1b*-deficient zebrafish while *col1a1b* (**Figure 2C**) expression levels remained stable. In addition, downregulation of *elnb* expression (**Figure 2A**) occurred in both mutant zebrafish lines.

**Figure 2:**
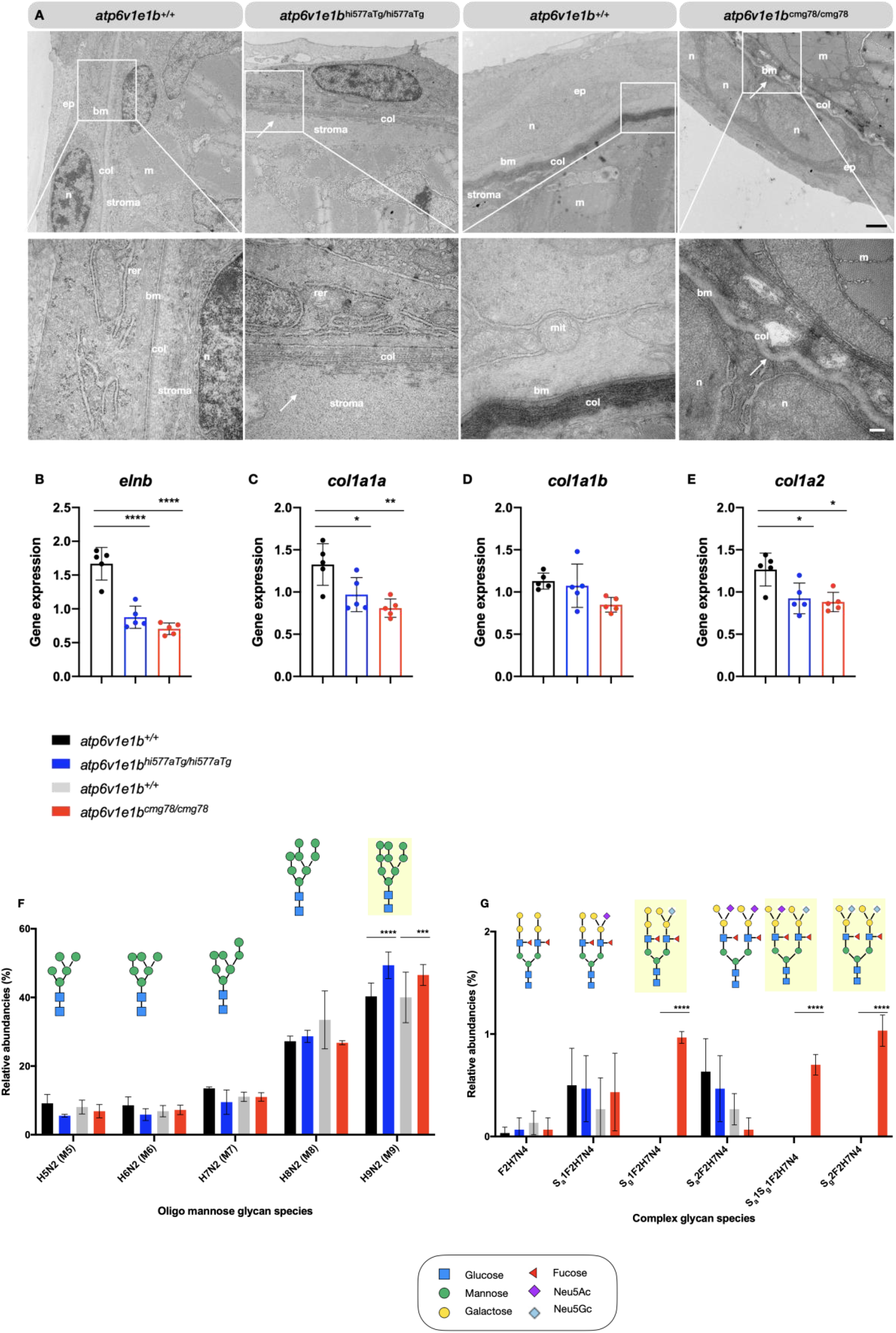
Profound epidermal and N-glycosylation alterations in *atp6v1e1b*-deficient zebrafish. (A) Transversal ultrathin section of the skin from WT and *atp6v1e1b*-deficient zebrafish at 4 dpf. A two-layered epidermis that is separated from the collagenous stroma of the dermis by a well-defined basement membrane (bm) is present at this developmental timepoint. In *atp6v1e1b*-deficient larvae, we observed a larger and disorganized collagenous stroma of the dermis (*atp6v1e1b*^hi577aTg/hi577aTg^) or coiled bm (*atp6v1e1b*^cmg78/cmg78^). Reproducible results were obtained in three independent experiments. Scale bar = 1 μm (top panel), scale bar = 200 nm (bottom panel). Ep: epidermis; bm: basement membrane; n: nucleus; col: collagenous fibrils; stroma: primary dermal stroma; m: muscle cell surface; rer: rough endoplasmic reticulum, mit: mitochondria. (B) Quantitative RT-qPCR analysis showed a significant decrease in gene expression of *elnb, col1a1a*, and *col1a2* in *atp6v1e1b*-deficient larvae at 4 dpf. Gene expression of *col1a1b* is similar in *atp6v1e1b*-deficient larvae compared to WT. Note that collagen type I is the most abundant collagen protein in the zebrafish skin. (F) Quantification of relative abundancies of MALDI-TOF-MS signals for oligomannose N-glycan species detected in deyolked lysates showed a significantly increased level of proteins containing 9 mannose residues in *atp6v1e1b*-deficient zebrafish. (G) Quantification of relative abundancies of MALDI-TOF-MS signals for complex N-glycan species detected in deyolked lysates demonstrated a significant increase for multiple complex N-glycan residues in *atp6v1e1b*cmg78/cmg78. Data are presented as mean ± SD from 3 biological replicates in G and F. 2-way ANOVA with Tukey test for multiple comparison. Yellow shading indicates differences between WT and *atp6v1e1b*-deficient larvae. Symbols represent monosaccharide residues. Graphical representation is based on the accepted convention from the Symbol Nomenclature for Glycans (Consortium for Functional Glycomics). Yellow circle, Gal; green circle, Man; red triangle, Fuc; purple diamond. Neu5Ac; light blue diamond, Neu5Gc. NeuAc: N-acetylneuraminic acid; NeuGc: N-glycolylneuraminic acid.

Since humans with ATP6V1E1 pathogenic variants show abnormal glycosylation, we investigated relative abundancies of N-glycans sugar chains in whole lysate proteins of *atp6v1e1b*-deficient zebrafish at 3 dpf [10]. We confirmed that the N-glycome profiles in whole zebrafish larval lysates were dominated by oligo-mannose glycans (M5 – M9) and had minor abundancies of complex glycans [36–38]. Complex glycans were exclusively composed of bi-antennary glycans without core fucosylation and antennary structures were of a uniform type characterized by mono- or disialylated Gal1-3Gal1-4(Fuc1-3)GlcNAc units according to MALDI-MS/MS. There was a significant increase in abundancy of oligo-mannose glycans, more specifically containing 9 mannose residues in *atp6v1e1b*-deficient larvae compared to wild-type (WT) controls (**Figure 2F**). We also observed a small reciprocal increase in complex glycans in *atp6v1e1b*^cmg78/cmg78^ larvae compared to WT controls (**Figure 2G**). *Atp6v1e1b*^cmg78/cmg78^ contained preferentially NeuGc, while *atp6v1e1b*^hi577aTg/hi577aTg^ exclusively had NeuAc as sialic acid species.

### Loss of *atp6v1e1b* does not induce a hypoxia-mediated response *in vivo*

*Atp6v1e1b*-deficient zebrafish show increased protein expression of the early endosomal markers early endosome antigen 1 (EEA1) and small rab GTPases 5 (Rab5) compared with WT controls at 4 dpf. The late endosomal marker small rab GTPase 7 (Rab7) and the lysosomal glycoprotein marker, LAMP1, on the other hand tended to show lower protein expression levels in *atp6v1e1b*-deficient zebrafish (**Supplemental Figure 2A-F**). Endosomal and lysosomal organelles have reduced acidification upon *atp6v1e1b* depletion in zebrafish, as shown by LysoTracker staining (**Supplemental Figure 2G**). This indicates impaired maturation of endo(lyso)somal vesicles in *atp6v1e1b*-deficient zebrafish.

Previous *in vitro* work has highlighted that decreased endo(lysosomal) acidification leads to reduced cytosolic bioavailability of iron and subsequent induction of HIFα signaling. HIFα-target genes (*egln3*, *vegfaa*, *vegfab*, *slc2a1a*, *slc2a1b*, *pfkfb3*, *angptl4* and *pdk1*) are not upregulated in *atp6v1e1b*-deficient zebrafish at 3 dpf (**Figure 3A-H**). Moreover, E3 medium supplemented with Ferric (Fe^3+^) ammonium citrate (FAC), which bypasses the endo(lyso)somal pathway through iron transporters in the plasma membrane [39], did not ameliorate the survival of *atp6v1e1b*-deficient zebrafish (**Figure 3I**). Also, hematopoiesis was not altered in *atp6v1e1b*-deficient zebrafish, as shown by o-dianisidine staining, providing further evidence of a normal availability of transferrin, hemoglobin and iron (**Figure 3J**).

**Figure 3:**
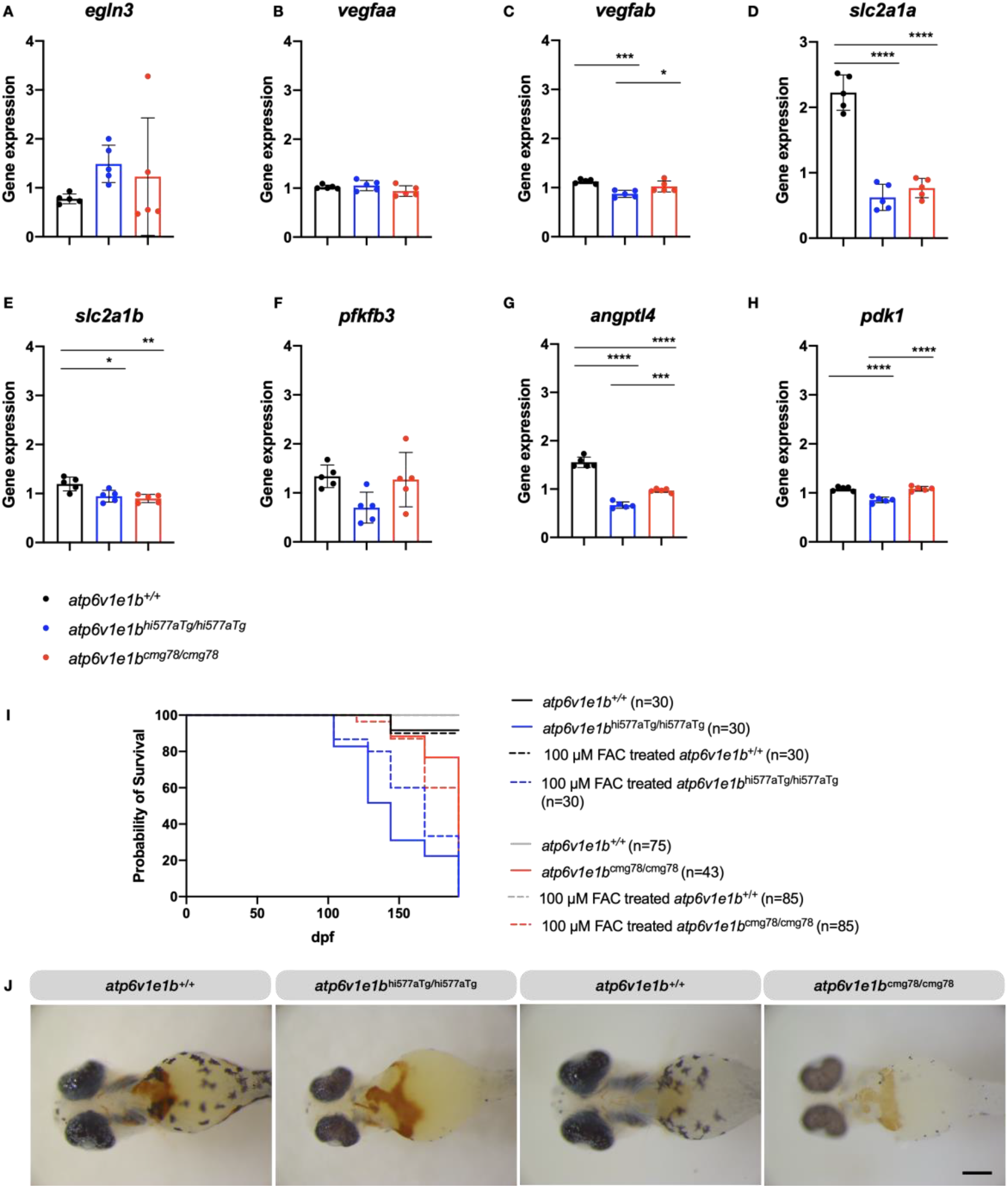
*In vivo* loss of *atp6v1e1b* does not induce a hypoxia-mediated response secondary to defective iron availability. (A-H) HIF1α-target genes are not upregulated at 3 dpf in *atp6v1e1b*-deficient zebrafish. No difference in gene expression of *egln3*, *vegfaa* and *pfkfb3* were found in *atp6v1e1b*-deficient zebrafish. Interestingly, the gene expression of *vegfab*, *slc2a1a*, *slc2a1b*, *angptl4* and *pdk1* was downregulated in *atp6v1e1b*-deficient zebrafish based on RT-qPCR. (J) Supplementation of 100 μM iron (Fe^3+^) ammonium citrate (FAC) does not improve survival of *atp6v1e1b*-deficient zebrafish. Kaplan-Meier curves for survival of *atp6v1e1b*-deficient zebrafish with and without FAC treatment. (J) A*tp6v1e1b*-deficient and WT zebrafish at 3 dpf exhibited normal levels of hemoglobin upon o-dianisidine staining. *Atp6v1e1b*^hi577aTg/hi577aTg^ (n=15) and their respective WT (n=15), *atp6v1e1b*^cmg78/cmg78^ (n=15) and their respective WT (n=15). Scale bar: 200 μM.

### Complex transcriptomic and metabolomic signature of *atp6v1e1b in vivo*

We performed an unbiased assessment of the transcriptional and metabolic response to the loss of *atp6v1e1b* at 3 dpf, before the gross morphological phenotype becomes apparent, in order to attempt to identify the primary mechanisms underlying the physiological manifestations of reduced intracellular acidification. Analysis of the transcriptomic data showed 112 significantly upregulated differentially expressed genes (DEGs) and 35 significantly downregulated DEGs in *atp6v1e1b*^hi577aTg/hi577aTg^ (**Figure 4A**). In addition, in *atp6v1e1b*^cmg78/cmg78^ 139 significantly upregulated DEGs and 29 significantly downregulated DEGs were found (**Figure 4B**). The top 30 most DEGs of *atp6v1e1b*-deficient zebrafish compared to WT controls are plotted in **Figure 4C**. We validated the expression levels of 10 genes picked from the top 150 most DEGs (**Supplemental Figure 3A**). Generally applicable gene-set enrichment (GAGE) for pathway analysis using the KEGG pathway database identified several altered pathways in *atp6v1e1b*^hi577aTg/hi577aTg^ and *atp6v1e1b*^cmg78/cmg78^ highlighting oxidative phosphorylation, fatty acid elongation, carbon metabolism, and steroid biosynthesis (**Figure 4D**). Several genes in the top 30 most DEGs are linked to GAGE pathway analysis, reactome pathway analysis and the observed phenotype (**Supplemental Figure 3B-C**). We observed significant upregulation of hatching enzyme 1 (*he1a*) and beta embryonic 1.1 hemoglobin (*hbbe1.1*). Moreover, we observed significant downregulation of beta embryonic 1.3 hemoglobin (*hbbe1.3*), alpha embryonic 1.1 hemoglobin (*hbae1.1*) and keratin 17 (*krt17*). Notch, Wnt and mTOR1 target genes were detected in our transcriptome dataset but the fold change differences were negligible.

**Figure 4:**
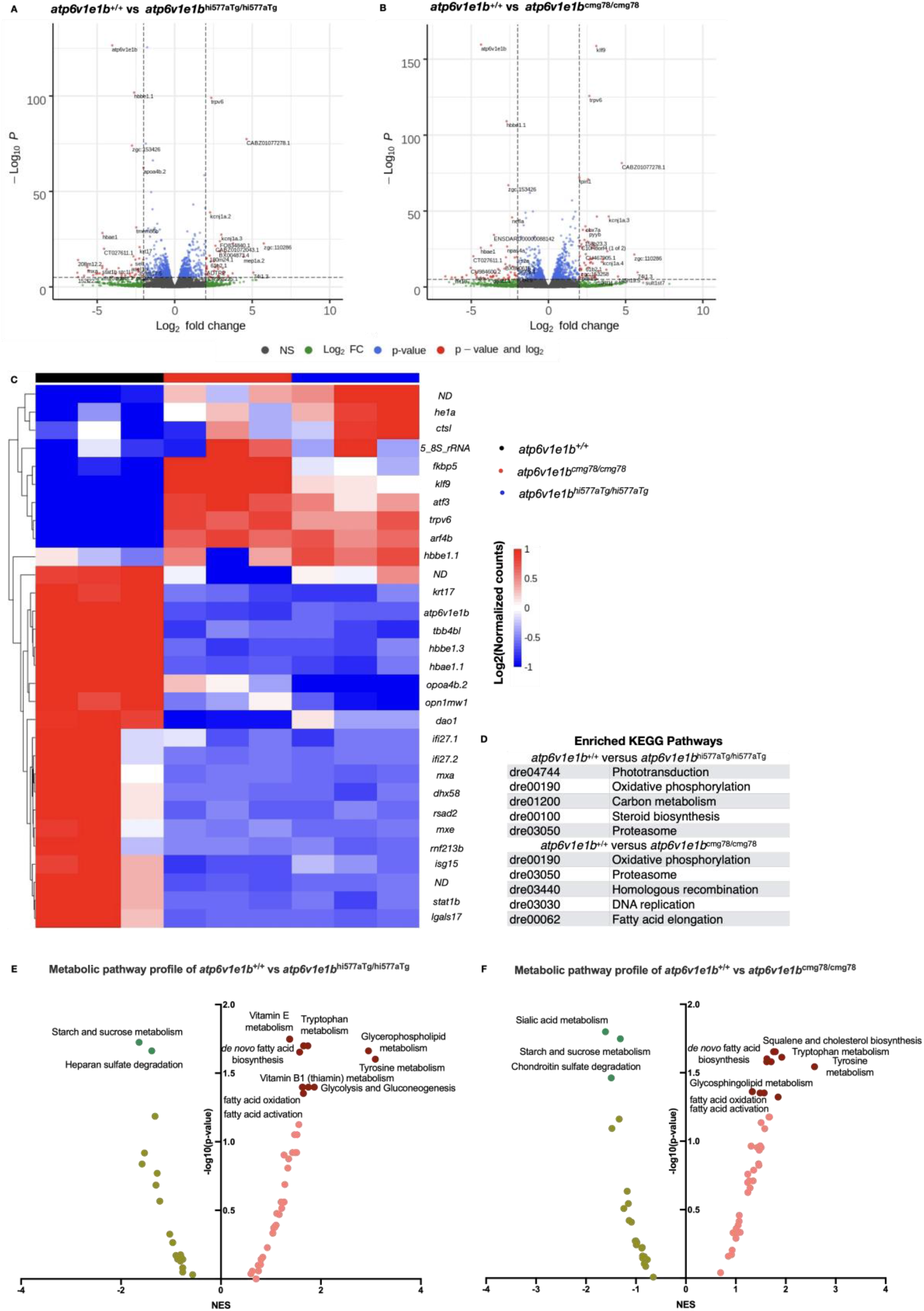
Transcriptomic and metabolomic signature of *atp6v1e1b* deficiency *in vivo*. (A) Volcano plot of RNA-sequencing data of *atp6v1e1b*^hi577aTg/hi577aTg^. (B) Volcano plot of RNA-sequencing data of *atp6v1e1b*^cmg78/cmg78^. The data represents overlapping genes from both ZFIN and Ensembl reference genome mapping (GRCz11). Red spots represent differentially expressed genes (DEGs). The horizontal line denotes the significance threshold (adjusted p<0.05) for DEGs. The vertical line denotes the Log2 fold change (Fc) threshold of 2. (C) Hierarchical clustering of the top 30 most DEGs from whole-body samples of *atp6v1e1b*-deficient and WT zebrafish at 3 dpf. DEGs are annotated with gene names of which the protein function is known. DEGs with unknown gene names are listed in **Supplementary Table 1**. Colors range from red (high expression) to blue (low expression). (D) Results of the KEGG pathway analysis showing the most enriched pathways in the differentially expressed gene list of the whole-body samples from the *atp6v1e1b*-deficient and WT zebrafish dataset. (E-F) Plots of an integrated analysis based on Metaboanalyst software (pathway tool) for a simplified view of contributing pathways in whole-body samples of *atp6v1e1b*-deficient zebrafish at 3 dpf. Full analysis based on GSEA parameters and Mummichog parameters are show in **Supplementary Table 2**. Dark green and brown symbols represent significantly enriched pathways. ND: not determined; NES: normalized enrichment score.

To complement the gene expression studies, we assessed the metabolomic profiles of *atp6v1e1b*-deficient zebrafish in order to uncover non-genomic changes in enzymatic activity of a range of cellular and metabolic processes. From an untargeted metabolic screen, we evaluated pathway-level enrichments based on significant m/z peaks. GSEA metabolic pathway activity profiles are plotted in **Figure 4E** and **Figure 4F** highlighting positive associations for the lipid metabolism, fatty acid metabolism, glycolysis and gluconeogenesis, and negative associations for the glycosaminoglycans. Though identification of a given peak based on its mass alone is challenging, we found two compounds significantly downregulated that, based on the accurate mass, could correspond to xanthopterin and hydroxymethylpterin. These metabolites are related to pigmentation which is absent in *atp6v1e1b*-deficient zebrafish. Our transcriptome and metabolome data show that multiple pathways are affected, particularly with prominent alterations in several parts of the lipid metabolism and oxygen consumption.

### Significant upregulation of sphingolipids, phospholipids and dihydroceramide upon *atp6v1e1b*-deficiency in zebrafish

We found a striking accumulation of electron-dense vesicular bodies in the yolk of the mutant zebrafish (**Figure 5A**). These organelles were packed with membranous, lamellar lipid-like material. *Atp6v1e1b*^hi577aTg/hi577aTg^ contained larger membranous whorls than *atp6v1e1b*^cmg78/cmg78^, in which multiple smaller membranous lipid whorls appeared in near proximity to each other (**Figure 5A**). Together with our omics results, this encouraged us to study 19 different lipid classes using HILIC LC-MS/MS lipidome analysis to determine the consequences of *atp6v1e1b*-deficiency at 3 dpf. Phosphatidylcholine (PC), and its derivatives 1-alkyl,2-acylphosphatidylcholine (PC-O), 1-alkenyl,2-acylphosphatidylcholine (PC-P), lysophosphatidylcholine (LPC), as well as phosphatidylglycerol (PG), phosphatidylinositol (PI), and sphingomyelin (SM) were significantly increased in *atp6v1e1b*-deficient zebrafish compared to WT controls (**Figure 5B-G**). Ceramides (CER) were not significantly altered in *atp6v1e1b*-deficient zebrafish compared to the WT controls. In contrast, dihydroceramide (DCER), the precursor of CER via the *de novo* ceramide synthesis pathway located in the ER, is significantly upregulated. Hexosylceramide (HexCer) and lactosylceramide (LaxCer) were significantly decreased in *atp6v1e1b*-deficient zebrafish compared to WT controls (**Figure 5I-L**). Levels of phosphatidylethanolamine (PE) and the related 1-alkyl,2-acylphosphatidylethanolamines (PE-O), 1-alkenyl, 2-acylphosphatidylethanolamines (PE-P) and lysophosphatidylethanolamine (LPE) showed a trend to be upregulated in *atp6v1e1b*-deficient zebrafish compared to WT controls (**Supplementary Figure 4A-D**). Similarly, fatty acids such as triacylgycerides (TG), diacylglycerides (DG), cholesterol esters (CE), and cholesterol (Chol) showed some increases (**Supplementary Figure 4E-F**). Inhibiting ceramide synthase, an enzyme that generates DCER in the *de novo* biosynthesis pathway in the ER, by administering Fumonisin B1 to the E3 medium at 1 dpf did not ameliorate the survival of *atp6v1e1b*-deficient zebrafish. The same observation was made when the ceramide transport protein (CERT), involved in the non-vesicular transport of ceramide from the ER membranes to Golgi apparatus for further processing to sphingomyelins, was inhibited by HPA-12 (**Supplementary Figure 5A-C**). Also, nicotinic acid, an inhibitor for the cholesterol esterification was not able to induce beneficial effects. These data suggest that targeting only this pathway is not sufficient to rescue the pleiotropic consequences of *atp6v1e1b* deficiency *in vivo*.

**Figure 5:**
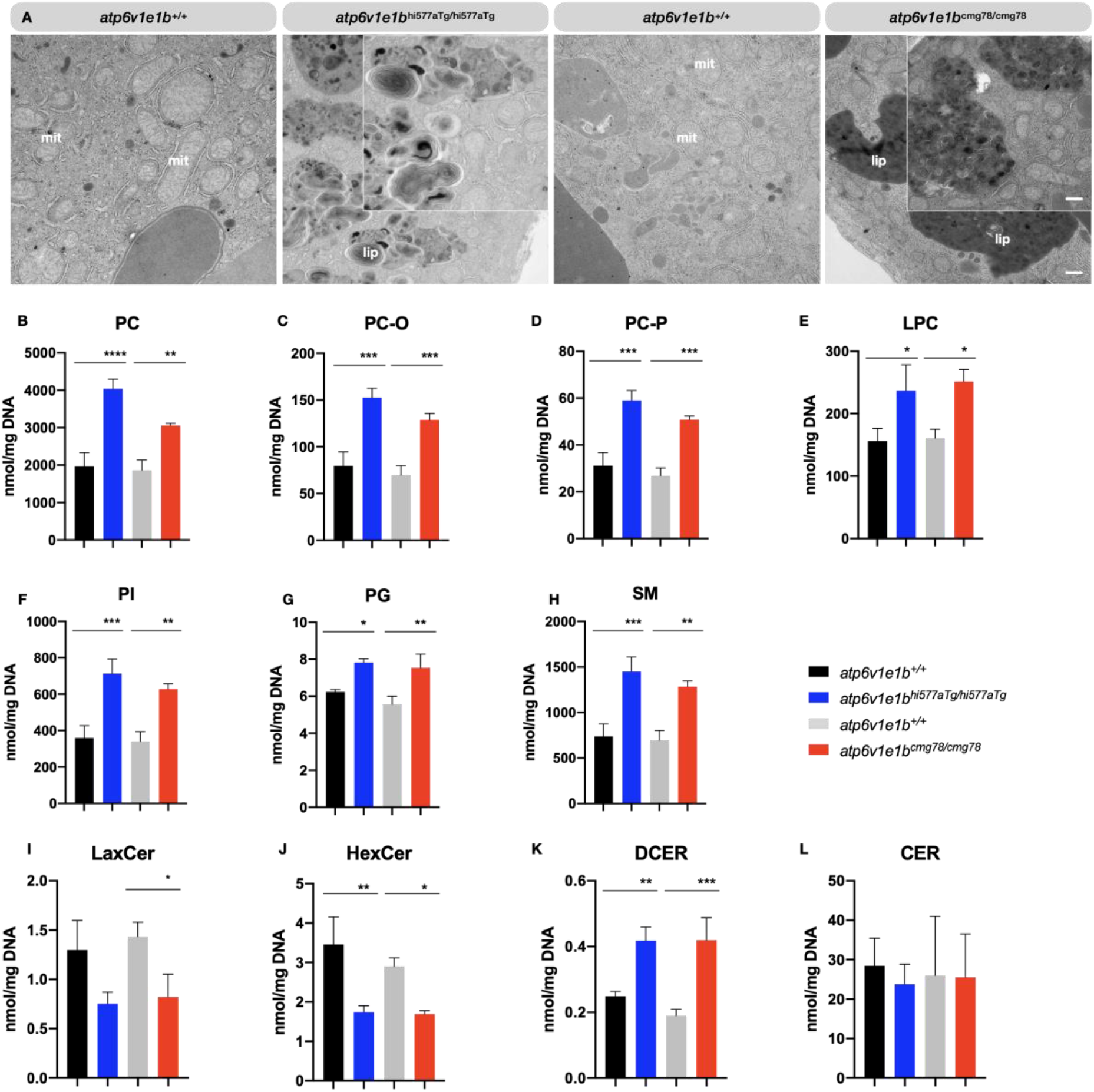
A *tp6v1e1b* depletion leads to higher amounts of sphingolipids and phospholipids in zebrafish larvae. (A) Representative images of ultrathin sections of the yolk from 4 dpf WT and *atp6v1e1b*-deficient zebrafish. *Atp6v1e1b*-deficient larvae reveal an accumulation of electron-dense vesicular bodies. Higher magnification of the lipid whorls is shown in the top right corner of the corresponding image. Scale bar = 500 nm (low magnification), scale bar = 200 nm (high magnification). Results are representative of three independent experiments. Mit: mitochondria; endo: endosomal derived multilamellar bodies; lip: lipid whorls. (B-L) HILIC LC-MS/MS lipidomic analysis demonstrates altered total levels of phospholipids, sphingolipids and sphingomyelins in 3 dpf *atp6v1e1b*-deficient zebrafish. Data are presented as mean ± SD from 3 biological replicates. PC: phosphatidylcholine; PC-O: 1-alkyl,2-acylphosphatidylcholine; PC-P: 1-alkenyl,2-acylphosphatidylcholine; LPC: lysophosphatidylcholine; PG: phosphatidylglycerol; PI: phosphatidylinositol; SM: sphingomyelin; CER: ceramides; DCER: dihydroceramides; HexCer: hexosylceramides; LacCer: lactosylceramides.

### Defective oxygen consumption in atp6v1e1b-deficient larvae

Upon TEM examination, we observed dilated mitochondria in *atp6v1e1b*^hi577aTg/hi577aTg^ and *atp6v1e1b*^cmg78/cmg78^ zebrafish. This, in combination with our transcriptome data suggests potential respiratory chain defects (**Figure 6A**), which prompted us to investigate the function of *atp6v1e1b*-deficient mitochondria. Oxygen consumption rate (OCR) measurements in WT controls, *atp6v1e1b*^hi577aTg/hi577aTg^ and *atp6v1e1b*^cmg78/cmg78^ zebrafish larvae at 2 dpf showed significantly decreased basal respiration in *atp6v1e1b*-deficient zebrafish compared to WT controls (**Figure 6B**). In addition, the basal extracellular acidification rate (ECAR) was lower in in *atp6v1e1b*-deficient zebrafish compared to WT controls **(Figure 6C**). Measurement of OCR and ECAR estimates the ATP generation from two different pathways: oxidative phosphorylation in the mitochondria and glycolysis in the cytosol [40]. Plotting ECAR against OCR of both mutant zebrafish and WT controls showed an altered metabolic state. *Atp6v1e1b*-deficient zebrafish showed a quiescent signature, while the corresponding WT controls are considered to be in a more energetic state (**Figure 6D**). Administration of the proton uncoupler carbonyl cyanide p-trifluoromethoxy-phenylhydrazone (FCCP) (**Figure 6F-I**) had no influence on the OCR in WT controls. FCCP however did increase the OCR in *atp6v1e1b*-deficient zebrafish, indicating spare respiratory capacity (**Figure 6F-I**). Administering oligomycin, a compound that inhibits the mitochondrial F-ATP synthase (complex V of the oxidative respiratory chain) [41], decreases the mitochondrial respiration which is linked to cellular ATP production in WT controls. Surprisingly however, we observed an immediate increase in OCR directly after administration of oligomycin in both *atp6v1e1b*-deficient zebrafish (**Figure 6E-G**). Considering the metabolic abnormalities in the *atp6v1e1b*-deficient zebrafish, we supplemented metabolites of the TCA cycle to the E3 medium of mutant larvae in order to attempt to ameliorate their phenotype. Administration of pyruvate, fumarate, oxaloacetic acid, succinate, glycolate, and L-(-)-acetic acid was not able to improve the survival of *atp6v1e1b*-deficient zebrafish (**Supplementary Figure 5D-I**). This indicates that targeting the TCA cycle alone is not sufficient to rescue the pleiotropic consequences of reduced *atp6v1e1b in vivo*.

**Figure 6:**
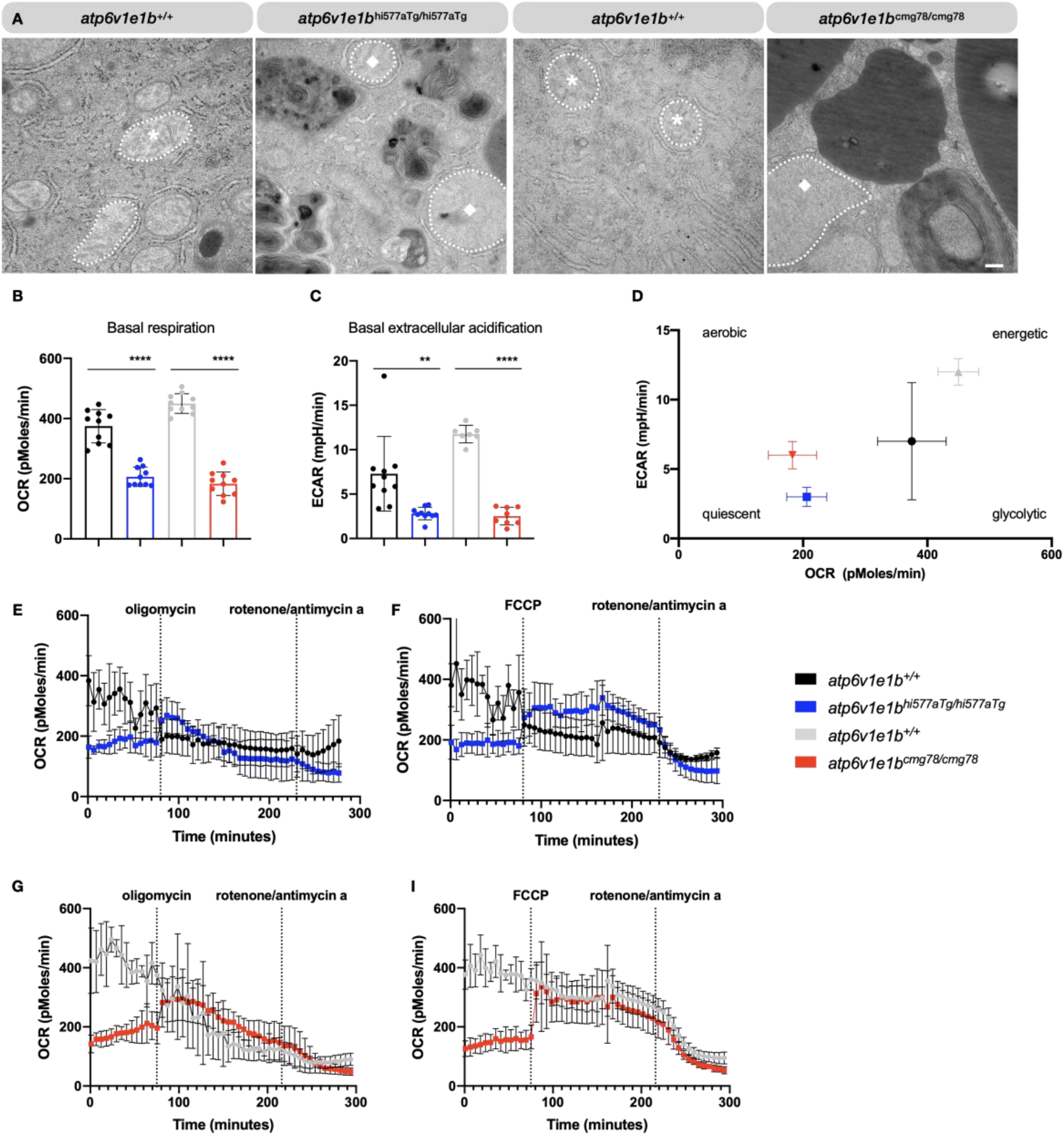
Early respiratory chain deficits resulting from *atp6v1e1b* depletion *in vivo.* (A) Representative images of ultrathin sections of the yolk from WT and *atp6v1e1b*-deficient zebrafish at 4 dpf, showing dilated mitochondria. Asterisk, normal mitochondria. Diamond, dilated mitochondria. Scale bar = 500 nm. Results are representative of three independent experiments. (B) The basal respiration rate and (C) extracellular acidification rate is significantly reduced in *atp6v1e1b*^hi577aTg/hi577aTg^ and *atp6v1e1b*^cmg78/cmg78^ zebrafish larvae at 2 dpf. (D) Plot of OCR vs. ECAR values showed that both mutant zebrafish larvae are in a more quiescent aerobic state than their WTs. (E-H) OCR in WT and *atp6v1e1b*-deficient zebrafish after administration of oligomycin and rotenone/antimycin a or FCCP and rotenone/antimycin a. OCR: oxygen consumption rate; ECAR: extracellular acidification rate.

## Discussion

Our data demonstrate that complete loss of *atp6v1e1b* in vivo mimics the ARCL type 2C phenotype [10]. *Atp6v1e1b*-deficiency was embryonically lethal in zebrafish, and resulted in craniofacial, cardiovascular, dermal and N-glycosylation abnormalities. It has previously been shown that increased N-glycans are associated with senescence and aging, and that luminal pH regulation is crucial for posttranslational modification in the Golgi compartment [42]. Indeed, our zebrafish model resulted in increased relative abundancies of different N-glycans as observed in ARCL type 2C patients [10].

We confirm that deficiency of v-ATPase plays a prominent role in the maturation of endo(lyso)somes [23]. It has been suggested that inhibition of v-ATPase function *in vitro* leads to intracellular Fe^2+^ deficits, due to impairment of the transferrin receptor pathway that requires low endosomal pH to release iron from the transferrin – transferrin receptor complex. Low cellular iron results in a stimulation of the HIF-mediated response, which could be reversed by iron supplementation *in vitro* [30–32]. We however found that HIF-target genes are not upregulated upon complete loss of *atp6v1e1b in vivo* and that supplementation of dietary FAC did not improve the survival rate. Under normal conditions the majority of iron is found as hemoglobin with supplemental iron stored intracellularly as ferritin, or bound to extracellular proteins [43]. Synthesis of heme occurs in the mitochondria and insufficient mitochondrial iron is known to decrease heme synthesis, resulting in lower hemoglobin levels [44] and hypochromic anemia [45, 46]. However, hemoglobin levels are normal in *atp6v1e1b*-deficient zebrafish suggesting normal levels of mitochondrial iron and that other mechanisms may bypass the impaired transferrin receptor pathway *in vivo*. One of the possibilities is that iron is delivered to mitochondria through siderophore-like molecules present in mammalian cells [46–48].

To understand the effects of reduced acidification *in vivo*, we performed an unbiased analysis of the transcriptome and metabolome in *atp6v1e1b*-depleted zebrafish at a developmental stage before gross morphological alterations became apparent, thereby reducing signals from secondary effects. Although similar hemoglobin levels were observed between *atp6v1e1b*-deficient zebrafish and WT controls, several hemoglobin genes (*hbbe1.3*, *hbae1.1*, and *hbbe1.1*) were regulated differently. Also, the reactome pathway analysis suggested oxygen transport deficits. Furthermore, KEGG pathway analysis on differentially regulated genes in *atp6v1e1b*-deficient zebrafish indicated enrichment of genes involved in oxidative phosphorylation, fatty acid elongation, carbon metabolism, and steroid biosynthesis pathways were enriched in *atp6v1e1b*-deficient zebrafish. It has been shown, mainly *in vitro*, that reduced acidification affects Notch, Wnt, and mTOR1 signaling [23]. However, major genes of these signaling pathways did not appear in the list of DEGs with Fc > 1. Since we studied the whole-body transcriptome of *atp6v1e1b*-deficient zebrafish larvae, it is nevertheless possible that effects limited to specific cell types are diluted in our data analysis.

Intriguingly, we found lipid-rich multilamellar bodies, characterized as lipid whorls, in the yolk of *atp6v1e1b*-deficient zebrafish. Lipid whorls are also observed in primary and secondary lysosomal storage disorders such as Niemann–Pick disease type C (NPC), which are characterized by reduced substrate clearance, and an accumulation of endo(lyso)somes [49, 50]. In NPC, deficiency of the NPC1 or NPC2 protein impairs the transmembrane transport of lipids, especially cholesterol, in the late endo(lyso)somal pathway [51]. It has previously been shown that v-ATPase subunits are associated with detergent resistant membranes present in late endosomes containing raft-like domains which are rich in cholesterol and sphingomyelin [52–54]. Our data confirmed a strong increase of PC levels and its derivatives, PG, PI, and SM levels in our *atp6v1e1b*-deficient zebrafish. Disturbances in sphingolipid metabolism have previously been linked to skeletal dysplasias, as well as cardiovascular and metabolic diseases [55–57]. In addition, unbiased analysis of the metabolic profile of *atp6v1e1b*-deficient zebrafish indicated glycosphingolipid metabolism, glycerophospholipid metabolism, glycolysis and gluconeogenesis, and fatty acid metabolism as key affected pathways. Taken together, our systems biology approach has shown that perturbation of v-ATPase function *in vivo* results in complex changes in multiple biological pathways, including phospholipid metabolism, sphingolipid metabolism, fatty acid metabolism, and oxygen transport deficits.

The results from our transcriptome analysis, highlighting alterations involved in oxygen transport and oxidative phosphorylation, together with the observation of dilated mitochondria, prompted us to further investigate the mitochondrial respiratory chain in our *atp6v1e1b* mutant models. While assessing mitochondrial respiration *in vivo*, we found that the uncoupling agent FCCP did not increase OCR capacity in 2 dpf WT larvae beyond baseline levels, suggesting that mitochondria are already at a maximal respiration rate at this stage of development. However, in *atp6v1e1b*-deficient zebrafish, which already have significantly reduced baseline respiration levels, FCCP did increase the OCR to similar levels as the observed baseline in WT larvae, suggesting suboptimal functioning of the respiratory chain in the mutants. Furthermore, after administering oligomycin, an ATP synthase inhibitor, we observed an unexpected increase in the OCR of *atp6v1e1b*-deficient zebrafish. This could result from a reverse direction of operation of the mitochondrial ATP synthase at baseline in *atp6v1e1b*-deficient zebrafish, a mechanism previously observed in mitochondrial dysfunction resulting in a disrupted electrochemical membrane potential [58, 59] and impaired mitochondrial ATP production. Given the evolutionary relationship between v-ATPase and F-ATP synthase, it is tempting to hypothesize that alteration of the E1 subunit of v-ATPase directly impacts F-ATP synthase function. A similar effect has already been suggested previously based on experiments in trypanosomes, where v-ATPase depletion was shown to affect F-ATP synthase coupling [60]. More experiments will need to be performed to better understand the functional interrelationship between these enzymes.

Considering the evidence gathered in our studies, we aimed to test whether modulating the identified pathways could ameliorate the phenotype in *atp6v1e1b*-deficient zebrafish. However, the administration of drugs interfering with the ceramide synthesis pathway or the TCA cycle did not improve the survival of *atp6v1e1b-*deficient zebrafish. These results suggest that we might have not yet found the right therapeutic target, or that compounds modulating multiple targets or pathways will be necessary to rescue the consequences of reduced organelle acidification caused by *atp6v1e1b* deficiency.

Plasma membrane v-ATPases promote cancer cell survival and malignancy since (i) they maintain a more alkaline intracellular environment that stimulates the Warburg effect and promotes actin polymerization near the v-ATPases enhancing cell mobility and (ii) they decrease the extracellular pH that favors activation of matrix metalloproteases and cathepsins, promoting the invasiveness and metastatic potential of cancer cells [17, 61–63]. Accordingly, *in vitro* experiments suggested that v-ATPase inhibition can be used as a viable therapeutic strategy to reduce cancer invasion and cell migration, and to avoid multi-drug resistance in cancer cells [21]. In light of these results, it has been proposed to consider the use of pan-v-ATPase inhibitors as a therapeutic treatment option for human melanoma [64]. Nevertheless, recent studies suggest that pan-v-ATPase inhibitors would likely be too toxic for therapeutic applications [1, 65], and it has been shown that prolonged exposure to specific v-ATPase inhibitors is lethal to mammalian cells [66–68]. Here, we provide additional evidence that genetic depletion of intracellular and plasma membrane v-ATPases causes lethality and disturbs multiple essential physiological processes *in vivo*. This urges the focus on the development of cancer cell specific v-ATPase inhibitors.

In summary, our mutant zebrafish models resemble the human ARCL type 2C phenotype with pleiotropic manifestations in multiple tissues. Our data suggest alternative routes, bypassing the transferrin receptor pathway, to deliver iron to mitochondria *in vivo*. Unbiased assessment of the transcriptional and metabolic response to the loss of *atp6v1e1b* identified the sphingolipid metabolism, fatty acid metabolism, and respiratory chain function as most prominently affected processes which we confirmed by analyzing the lipidome and the oxygen consumption in *atp6v1e1b*-deficient zebrafish. Pharmacological intervention focusing on one of these pathways however did not ameliorate the survival of *atp6v1e1b*-deficient zebrafish suggesting that the multisystemic biological effects of *atp6v1e1b* might require simultaneous targeting of multiple mechanisms to rescue the loss of *atp6v1e1b in vivo*. Overall, our study has identified new complex biological effects of *atp6v1e1b* deficiency *in vivo*, which deserve further investigation and might be leveraged to develop new therapeutics in the future.

## Materials & methods

### Zebrafish lines and maintenance

Zebrafish lines were housed in a Zebtec semi-closed recirculation housing system at a constant temperature (27– 28 °C), pH (~7.5), conductivity (~550 μS) and light/dark cycle (14/10). Fish were fed twice a day with dry food (Gemma Micro, Skretting) and once with artemia (Ocean Nutrition, Essen, Belgium). *Atp6v1e1b*^hi577aTg/+^ was purchased from the Zebrafish International Research Center (ZIRC, University of Oregon, Eugene, United States). *Atp6v1e1b*^hi577aTg/hi577aTg^ constitutes a retroviral insertion in the 5’ UTR of the gene [34, 69]. *Atp6v1e1b*^cmg78/cmg78^ was generated using CRISPR-Cas9 mutagenesis according to the workflow previously described [70]. Zebrafish were genotyped with primers listed in **Supplemental Table 3**. We adhered to the general guidelines, in agreement with EU Directive 2010/63/EU for laboratory animals, for zebrafish handling, mating, embryo collection and maintenance [71, 72]. Approval for this study was provided by the local committee on the Ethics of Animal Experiments (Ghent University Hospital, Ghent, Belgium; Permit Number: ECD 17/63K and ECD 18/05).

### Drug administration

Compounds were administered to zebrafish embryos at 1 dpf, after chorion removal in E3 embryo medium (containing 5 mM NaCl, 0.17 mM KCl, 0.33 mM CaCl_2_, 0.33 mM MgSO_4_, and 50 mM HEPES pH 7.1) in a Petri dish. In cases in which drugs were dissolved in DMSO dilutions were made so the final DMSO concentration did not exceed 1% in the Petri dish with the exception of nicotinic acid. Nicotinic acid was dissolved in NaOH. The final NaOH concentration did not exceed 0.03% in the Petri dish. The relevant vehicle control was used for each experiment. Drugs are listed in **Supplemental Table 4**.

### Evaluation heart function and vasculature

Video microscopy was performed on an Axio Observer.Z1 inverted microscope attached to a monochrome Axiocam 506 camera (Carl Zeiss Microscopy GmbH, Jena, Germany). Sequential images of the heart and blood flow region of interest were obtained in lateral position at 50 frames/sec at 3 dpf. Videos of the heart were manually analyzed using Fiji software [73]. The short and long axis of an ellipse fitted to the outline of the outside ventricular wall was measured 3 times in systole and in diastole [74]. Based on the assumption that the ventricle of the zebrafish embryo is a prolate spheroid [75], the systolic and diastolic volume was calculated. Videos of the blood flow through the tail vasculature were analyzed as previously described [76].

### Evaluation of the vasculature structure

Zebrafish were outcrossed to the *Tg(kdrl:eGFP*) zebrafish line in order to visualize endothelial cells. Zebrafish larvae of 5 dpf were embedded in 0.8% seaPlaque low melting Agarose (Lonza, Basel, Switzerland) supplemented with 160 mg/L tricaine (Sigma-Aldrich, Saint Louis, USA) in order to minimize the artefactual movements. Z-stacks were taken on an Axio Observer.Z1 inverted microscope attached to a monochrome Axiocam 506 camera (Carl Zeiss Microscopy GmbH) and analyzed using the stack focuser algorithm in Fiji software. The segmental area of the aorta between the first (AA3) and the second branchial arch (AA4) [77], diameter of aorta proximal to bulbus, average of 3 segments between branches of the ventral aorta and the diameter of the bulbus arteriosus were measured.

### Transmission electron microscopy

Zebrafish mutant larvae of 4 dpf were fixed and processed for ultrastructural analysis as previously described [78]. Sections were viewed with Jeol JEM 1010 TEM (Jeol Ltd., Tokyo, Japan) equipped with a CCD side mounted Veleta camera operating at 60 kV. Experiments were performed in collaboration with the TEM facility of the Nematology Research Unit. Pictures were digitized using a Ditabis system (Ditabis Ltd., Pforzheim, Germany).

### Assay of motor function, survival, Alcian Blue and lysosomal staining

Touch-evoked escape response was measured at 3 dpf using a scale from 1 to 4: 1, no movement; 2, local muscle contraction of the embryo; 3, short distributed swim movement; 4, normal swim movement towards the edge of the Petri dish. Zebrafish were touched with the tip of a P10 pipette at the end of their tail. Zebrafish mortality was scored based on cardiac arrest and tissue degradation. Survival analysis was plotted using Kaplan-Meier curves. Cartilage patterns were stained with Alcian Blue as previously described [79]. Craniofacial structures were measured with Fiji software and normalized to the length of the head. Stained specimens were analyzed with a Leica M165 FC Fluorescent Stereo Microscope (Leica Microsystems, GmbH, Wetzlar, Germany). For lysosomal imaging, zebrafish larvae at 3 dpf were incubated with LysoTracker Red DND-99 (Life Technologies) which contains fluorescent acidotropic probes for labeling and tracking of acidic organelles. Zebrafish larvae were subsequently washed and were embedded in 0.8% seaPlaque low melting Agarose (Lonza) supplemented with 160 mg/L tricaine (Sigma-Aldrich). Brain area was imaged using a Leica TCS LSI zoom confocal microscope.

### Western blot

For western blot analysis, 20 zebrafish larvae were homogenized with a pestle in SDS Laemlli buffer (10% glycerol, 2% SDS, 0.5M Tris HCl pH 8,0 and dH_2_O) at 4 dpf. Protein samples were subjected to 7%, 10% or 15% SDS polyacrylamide gel electrophoresis, and transferred to a polyvinyldifluoride (PVDF) or nitrocellulose (NC) membrane (Thermo Fisher Scientific, Waltham, Massachusetts, United States) with the iBlot 2 Dry Blotting System (Thermo Fisher Scientific). Imaging was performed with an Amersham Imager 600 CCD camera (GE Healthcare, Chicago, Illinois, United States) and analyzed using Fiji software. The following primary antibodies were used: monoclonal rabbit anti-Rab5 antibody (1:1000, C8B1, Cell Signaling Technology, Danvers, Massachusetts, United States), monoclonal rabbit anti-Rab7 antibody (1:1000, D95F2, Cell Signaling Technology), monoclonal rabbit anti-EEA1 antibody (1:1000, C45B10, Cell Signaling Technology) and polyclonal rabbit anti-LAMP1 (1:1000, ab24170, Abcam, Cambridge, United Kingdom). HRP-conjugated goat anti-rabbit IgG was used as a secondary antibody (1:2000, #7074, Cell Signaling Technology).

### Oxygen measurements

An XF24 Extracellular Flux Analyzer (Agilent Technologies, Santa Clara, CA, USA) was used to measure oxygen consumption rate (OCR) and extracellular acidification rate (ECAR). 10 of the 24 wells on an islet capture microplate (Agilent Technologies) were used for mutant zebrafish larvae, 10 wells were used for their respective WT controls and 4 wells served as quality control (QC). Four zebrafish larvae were put together in 1 well. Zebrafish larvae were incubated in 1x E3-medium buffered with 4mM 4-(2-hydroxyethyl)-1-piperazineethanesulfonic acid (HEPES) (Life Technologies, Carlsbad, CA, USA). Seahorse XF Cell Mito Stress kit (Agilent Technologies) was used according to manufacturer’s instructions and modified for zebrafish purposes as described in previous publications [80–82]. OCR data and ECAR data were exported using Wave Desktop Software (Agilent Technologies).

### Lipid extraction, mass spectrometry and data analysis

40 larvae of 3 dpf were homogenized in 500 μl H_2_O with the Precellys system and a sample volume equal to 10 μg of DNA was diluted in 700 μl H_2_O and mixed with 800 μl 1 N HCl:CH3OH 1:8 (v/v), 900 μl CHCl_3_ and 200 μg/ml of the antioxidant 2,6-di-tert-butyl-4-methylphenol (BHT) (Sigma Aldrich). 3 μl of SPLASH^®^ LIPIDOMIX^®^ Mass Spec Standard, (Avanti Polar Lipids, Alabaster, AL, USA) was spiked into the extraction mix. The organic fraction was evaporated using a Savant Speedvac spd111v (Thermo Fisher Scientific) at room temperature and the remaining lipid pellet was stored at −20°C under argon. Just before mass spectrometry analysis, lipid pellets were reconstituted in 100% ethanol. Lipid species were analyzed by liquid chromatography electrospray ionization tandem mass spectrometry (LC-ESI/MS/MS) on a Nexera X2 UHPLC system (Shimadzu, Kioto, Japan) coupled with hybrid triple quadrupole/linear ion trap mass spectrometer (6500+ QTRAP system; AB SCIEX). Chromatographic separation was performed on a XBridge amide column (150 mm × 4.6 mm, 3.5 μm; Waters) maintained at 35°C using mobile phase A [1 mM ammonium acetate in water-acetonitrile 5:95 (v/v)] and mobile phase B [1 mM ammonium acetate in water-acetonitrile 50:50 (v/v)] in the following gradient: (0-6 min: 0% B → 6% B; 6-10 min: 6% B → 25% B; 10-11 min: 25% B → 98% B; 11-13 min: 98% B → 100% B; 13-19 min: 100% B; 19-24 min: 0% B) at a flow rate of 0.7 mL/min which was increased to 1.5 mL/min from 13 minutes onwards. SM, CE, CER, DCER, HCER, LCER were measured in positive ion mode with a precursor scan of 184.1, 369.4, 264.4, 266.4, 264.4 and 264.4 respectively. TAG, DAG and MAG were measured in positive ion mode with a neutral loss scan for one of the fatty acyl moieties. PC, LPC, PE, LPE, PG, LPG, PI, and LPI were measured in negative ion mode with a neutral loss scan for the fatty acyl moieties. Lipid quantification was performed by scheduled multiple reactions monitoring (MRM), the transitions being based on the neutral losses or the typical product ions as described above. Peak integration was performed with the MultiQuantTM software version 3.0.3. Lipid species signals were corrected for isotopic contributions (calculated with Python Molmass 2019.1.1) and were normalized to internal standard signals. Unpaired T-test p-values and FDR corrected p-values (using the Benjamini/Hochberg procedure) were calculated in Python StatsModels version 0.10.1.

### Transcriptomics

Total RNA was isolated from whole zebrafish larvae at 3 dpf from *atp6v1e1b*^hi577aTg/hi577aTg^ and *atp6v1e1b*^cmg78/cmg78^ zebrafish together with their respective WT controls as described previously [83]. RNA integrity was checked by 2100 Bioanalyzer (Agilent Technologies). 10 zebrafish larvae of each genotype were pooled in 1 sample to obtain sufficient heterogeneity amongst samples. A sequencing library was prepared using the TruSeq^®^ Stranded mRNA Library Prep (Illumina, San Diego, CA, USA) supplemented with TruSeq^®^ RNA Single Indexes Set A (Illumina), according to the manufacturer’s instructions. Paired-end sequencing was performed on an HiSeq 3000 sequencer (Illumina) with the HiSeq 3000/4000 SBS kit (150 cycles) according to the manufacturer’s protocol. Data analysis was done in collaboration with the bio-informatics core (Department of Biomolecular Medicine, UGent). An RNA-seq pipeline was used that was published by the nf-core community. This pipeline was executed using the Nextflow [84] engine for computational workflows and comprises several processing steps [85]. QC analysis of the RNA-seq data was performed with FastQC and MultiQC [86]. TrimGalore was used to remove adapter contamination and to trim low-quality regions. Duplicate reads were identified with MarkDuplicates (Picard). Subsequently, all cleaned and trimmed reads that passed QC were aligned to GRCz10 using STAR aligner [87]. The index trimmed paired-end 150 base pair reads were aligned to the zebrafish GRCz10 reference genome. Gene counts were computed using the featureCounts package [88]. Differential expression analysis subsequently was performed on these gene counts using DESeq2 [89]. Differentially expressed genes were identified using a fold change cut-off >1 and FDR=0.05. Finally, GO enrichment, GO pathway (http://geneontology.org/) and KEGG pathway (https://www.genome.jp/kegg) analysis were performed on differentially expressed gene sets using the generally applicable gene-set enrichment for pathway analysis (GAGE) algorithm [90]. RNA sequencing data used for the gene expression analysis of both *atp6v1e1b-*deficient zebrafish and WT controls have been deposited in the ArrayExpress database at EMBL-EBI under accession number E-MTAB-8824, and can be accessed at the following link: https://www.ebi.ac.uk/arrayexpress/experiments/E-MTAB-8824. This link is still under embargo but can be requested at ArrayExpress for review.

### RT-qPCR

Quantitative reverse transcription PCR (RT-qPCR) was performed as described previously [83]. Total RNA was extracted in quintuplicate, in which 10 zebrafish larvae were pooled per sample. Assays were prepared with the addition of ssoAdvanced SYBR Green supermix (Bio-Rad Laboratories) and were subsequently run on a LightCycler^®^ 480 Instrument II (Roche, Basel, Switzerland). Primers were designed using Primer-BLAST (**Supplemental Table 5**). Biogazelle qBase+3.0 Software was used for data analysis using *bactin2*, *elfa* and *gapdh* for normalization.

### Metabolomics

In order to assess the effect of *atp6v1e1b* reduction on their metabolome, 3 batches of ∼100 *atp6v1e1b*^hi577aTg/hi577aTg^, *atp6v1e1b*^cmg78/cmg78^ and their WT controls were harvested at 3 dpf. Larvae were homogenized with a pestle in dH_2_O and the larvae were frozen until processed. Samples were subjected to Ultra Performance Liquid Chromatography High Resolution Mass Spectrometry (UPLC-HRMS) at the VIB Metabolomics Core Ghent (VIB-MCG). 10 ul was injected on a Waters Acquity UHPLC device connected to a Vion HDMS Q-TOF mass spectrometer Chromatographic separation was carried out on an ACQUITY UPLC BEH C18 (50 × 2.1 mm, 1.7 μm) column from Waters, and temperature was maintained at 40°C. A gradient of two buffers was used for separation: buffer A (99:1:0.1 water:acetonitrile:formic acid, pH 3) and buffer B (99:1:0.1 acetonitrile:water:formic acid, pH 3), as follows: 99% A for 0.1 min decreased to 50% A in 5 min, decreased to 30% from 5 to 7 minutes, and decreased to 0% from 7 to 10 minutes. The flow rate was set to 0. 5 mL min−1. Both positive and negative Electrospray Ionization (ESI) were applied to screen for a broad array of chemical classes of metabolites present in the samples. The LockSpray ion source was operated in positive/negative electrospray ionization mode under the following specific conditions: capillary voltage, 2.5 kV; reference capillary voltage, 2.5 kV; source temperature, 120°C; desolvation gas temperature, 600°C; desolvation gas flow, 1000 L h−1; and cone gas flow, 50 L h−1. The collision energy for full MS scan was set at 6 eV for low energy settings, for high energy settings (HDMSe) it was ramped from 28 to 70 eV. Mass range was set from 50 to 1000Da, scan time was set at 0.1s. Nitrogen (greater than 99.5%) was employed as desolvation and cone gas. Leucine-enkephalin (250 pg μL−1 solubilized in water:acetonitrile 1:1 [v/v], with 0.1% formic acid) was used for the lock mass calibration, with scanning every 1 min at a scan time of 0.1 s. Profile data was recorded through Unifi Workstation v2.0 (Waters). Data processing was performed with Progenesis QI software version 2.4 (Waters) for chromatogram alignment and compound ion detection. The detection limit was set at maximum sensitivity. In ESI^+^ and ESI^−^ ionization, 9795 and 8959 features were detected and aligned, respectively. Post-processing was done with the online MetaboAnalyst 4.0 tool (http://www.metaboanalyst.ca) [91, 92]. Briefly, the features were normalized by log transformation and Pareto scaling upon statistical analysis. T-test with FDR<0.1 identified 329 features for *atp6v1e1b*^hi577aTg/hi577aTg^ compared to WT controls and 521 features as significant for *atp6v1e1b*^cmg78/cmg78^ compared to WT controls. Functional interpretation directly from m/z peaks was performed based on a well-established GSEA algorithm [92]. Identifications of significant m/z peaks (one-way ANOVA, FDR<0.02) was attempted with Chemspider including following libraries: Chebi, Humann Metabolome database, Golm Metabolome database, The national compound collection, Targetmol, Sigma Aldrich, Pubmed, Natural products Atlas, NIST Spectra, NIST, Mcule, Massbank, KEGG, FoodDB, Extrasynthese and Analyticon Discovery.

### N-Glycomics

In order to assess the effect of *atp6v1e1b* reduction on N-linked glycosylation, 3 batches of ∼400 *atp6v1e1b*^hi577aTg/hi577aTg^, *atp6v1e1b*^cmg78/cmg78^, and WT controls were harvested at 3 dpf. Yolks were manually removed from one batch and the larvae were frozen until processed. Larvae were homogenized with a pestle in 1% octyl-glucoside (Life Technologies) in dH_2_O. Samples were concentrated by Amicon ultrafiltration (Merck Millipore, Burlington, MA, USA). Samples were subjected to trypsin and peptide-N-glycosidase treatment before solid-phase extraction on C18 cartridges. Samples were dried by vacuum rotation after which glycans were methylated. Methylated glycans were applied onto a MALDI stainless steel target (Bruker Daltonics, MA, USA) together with an equal volume of matrix (α-cyano-4-hydroxy-cinnamic acid, saturated solution in 50% acetonitrile/0.1% aqueous TFA). MALDI-MS analysis was performed in the positive ion reflectron mode (HV acceleration 25 kV) on an UltrafleXtreme MALDI-TOF-TOF mass spectrometer (Bruker Daltonics) using the acquisition software FlexControl 3.3 and the data evaluation software FlexAnalysis 3.3 (Bruker Daltonics). The mass range selected for detection was from 500 to 5.000 Da. In general, 5.000 laser shots were accumulated at 1 kHz acquisition for each sample spot.Partial sequencing of glycans by MALDI-MS/MS was performed by laser-induced dissociation (LID) in the post-source decay (PSD) mode. Mass annotation was performed in FlexAnalysis and peaklists were generated after manual inspection of the entire mass range for correct isotope peak annotation. To assist oligosaccharide identification on the basis of sugar composition, the automatic tool for sugar mass increments was used. The molecular masses M^+^Na, M^+^Na-32, and M^+^Na-54 were compared with standard mass lists, however a final identification was based on combined MS and MS/MS data. Fragmentation patterns were evaluated manually using fragmentation ion masses predicted by the GlycoWorkbench software tool.

### Quantitative analysis and statistics

Data processing and statistical analyses were performed using Graph Pad Prism version 8.01 (GraphPad Software, San Diego, CA, USA), which was used to generate each of the graphs shown in the figures, performing statistical tests as specified in figure legend or Materials and Methods. All data are presented as mean ± SD. * P-value < 0.05; ** P-value < 0.01; *** P-value < 0.001; **** P-value < 0.0001. One-way ANOVA with correction for multiple comparison using the Bonferroni method was used for the majority of the analysis, except if it is mentioned otherwise in the figure legend.

## Acknowledgements

We would like to thank Ms. Petra Vermassen, Ms. Hanna De Saffel, Ms. Zoë Malfait, and Ms. Myriam Claeys for their technical support. Electron microscopy was performed in the TEM facility of the Nematology Research Unit. Secondly, we would also like to thank Ms. Jelena Jovanovic for the practical assistance with the seahorse experiments. Lipidomics was performed by Lipometrix, core facility at the KU Leuven (www.lipometrix.be). The Ghent University Hospital is member of the European Reference Network (ERN)-Skin, VascERN, and ReCONNET.

## CRediT author statement

Lore Pottie: Conceptualization, Data Curation, Formal Analysis, Investigation, Methodology, Project Administration, Visualization, Writing – Original Draft Preparation, Writing – Review & Editing.

Wouter Van Gool: Data Curation, Formal Analysis, Software, Writing – Review & Editing.

Michiel Vanhooydonck: Investigation, Writing – Review & Editing.

Franz-Georg Hanisch: Investigation, Resources, Writing – Review & Editing.

Geert Goeminne: Investigation, Resources, Writing – Review & Editing.

Andreja Rajkovic: Resources, Writing – Review & Editing.

Paul Coucke: Funding Acquisition, Resources, Writing – Review & Editing.

Patrick Sips: Conceptualization, Supervision, Writing – Review & Editing.

Bert Callewaert: Conceptualization, Funding Acquisition, Resources, Supervision, Project Administration, Writing – Review & Editing.

## Supporting information

**Supplemental Figure 1:**
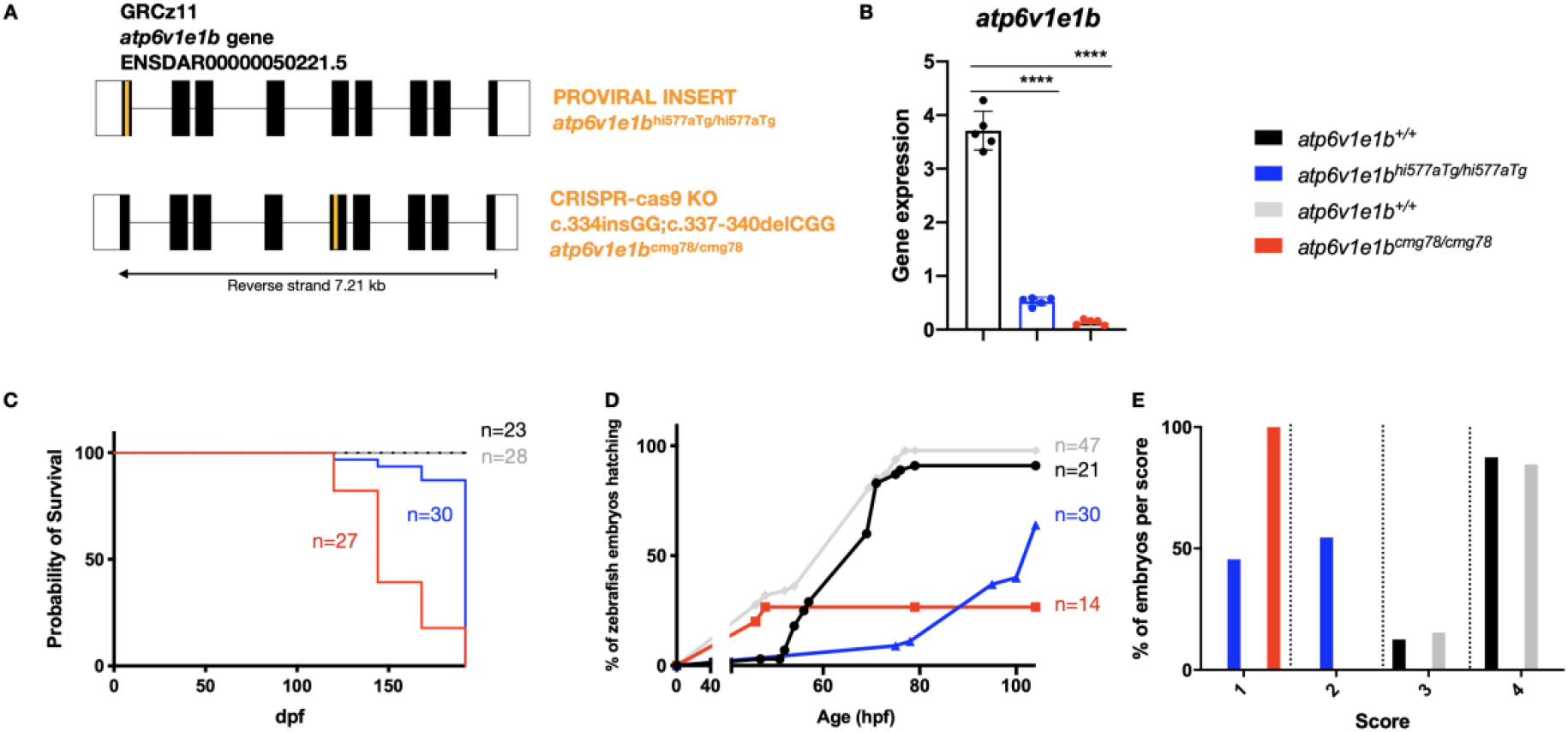
Knock-out (KO) of *atp6v1e1b* in zebrafish shows an early phenotypic read-out. (A) Schematic representation of the *atp6v1e1b* gene (transcript ENSDAR00000050221.5). The orange line represents the position of the proviral insert (*atp6v1e1b*^hi577aTg/+^) or the target site of the sgRNA-guided CRISPR-Cas9 indel mutagenesis (*atp6v1e1b*^cmg78/+^). (B) Quantitative reverse transcription PCR (RT-qPCR) analysis showed a significant decrease in expression of *atp6v1e1b* in both zebrafish models at 3 dpf. No significant difference could be detected in *atp6v1e1b* relative expression levels between *atp6v1e1b*^hi577aTg/hi577aTg^ and *atp6v1e1b*^cmg78/cmg78^. (C) Loss of a*tp6v1e1b* causes embryonic mortality in zebrafish. Kaplan-Meier curves for survival of *atp6v1e1b*-deficient zebrafish. (D) Hatching pattern of *atp6v1e1b*-deficient zebrafish. *Atp6v1e1b*-defcient zebrafish showed delayed or no spontaneous hatching (E) Touch-Evoked Escape test (TEE-test) of *atp6v1e1b*^hi577aTg/hi577aTg^ larvae (n=22) and their respective wild-type (WT) (n=8), *atp6v1e1b*^cmg78/cmg78^ (n=26) larvae and their respective WT (n=26). The reaction after touch with a thick blunt needle was scored as follows: (1) no movement, (2) local muscle contractions of the embryo, (3) short distance swim movement and (4) normal swim movement towards the edge of the Petri dish.

**Supplemental Figure 2:**
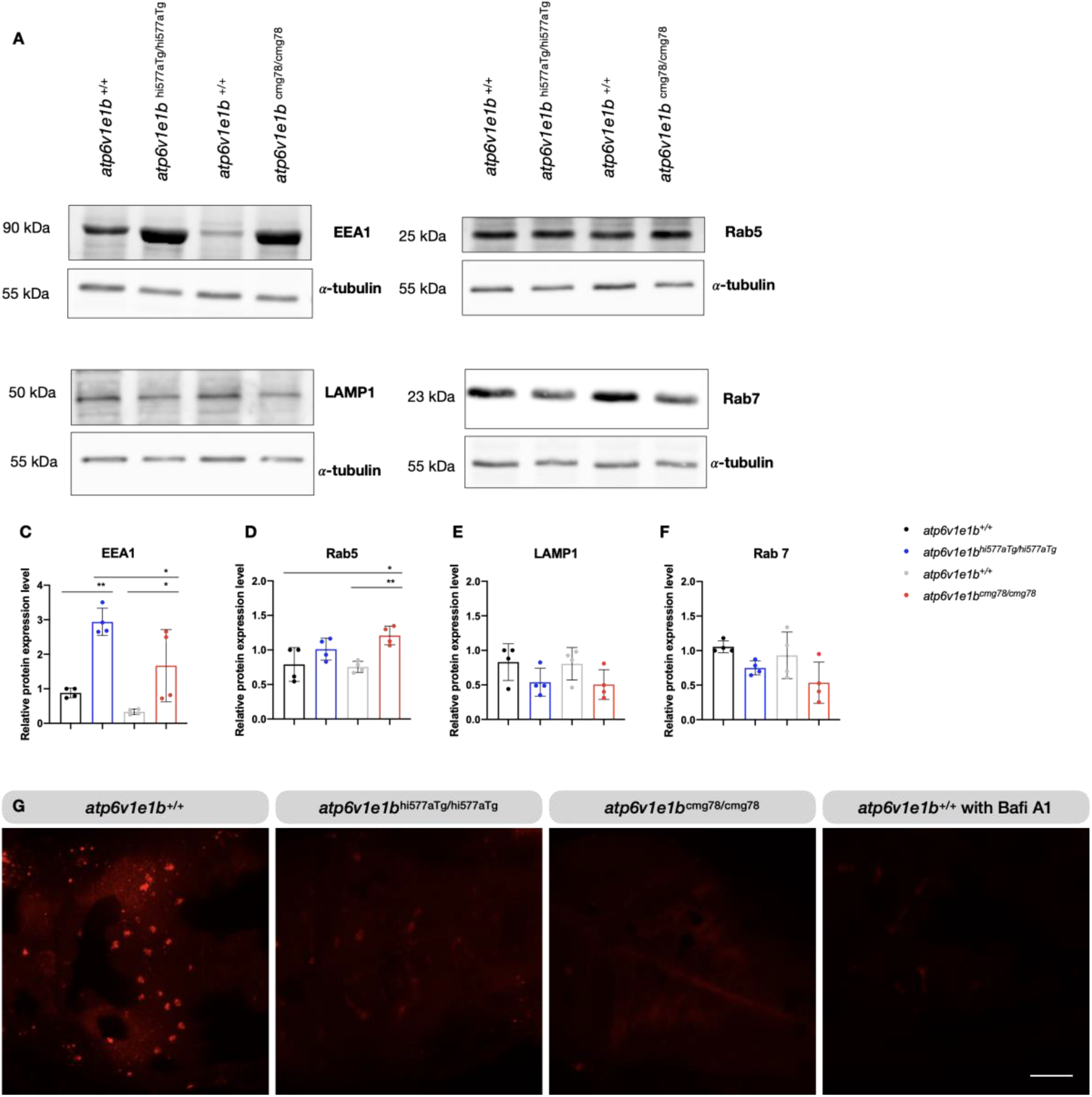
*Atp6v1e1b*-deficiency blocks the maturation of endosomal and lysosomal vesicles in zebrafish, retaining similar canonical function as in mammalian species. (A) Western Blot analysis of whole larvae of *atp6v1e1b*^hi577aTg/hi577aTg^ and *atp6v1e1b*^cmg78/cmg78^ and their corresponding WT. We confirmed equal loading by staining for α-tubulin. (C-F) Quantification of the relative protein levels of EEA1, Rab5, LAMP1 and Rab7 in whole zebrafish lysates, normalized to total α-tubulin levels. (G) Representative confocal images are shown. Acidified lysosomes are visualized in red. *Atp6v1e1b*^+/+^ (n=12), *atp6v1e1b*^hi577aTg/hi577aTg^ (n=12), *atp6v1e1b*^cmg78/cmg78^ (n=12) and *atp6v1e1b*^+/+^ treated with 1,6 μM Bafilomycin A1 (n=9). Scale bar= 100 μm.

**Supplemental Figure 3:**
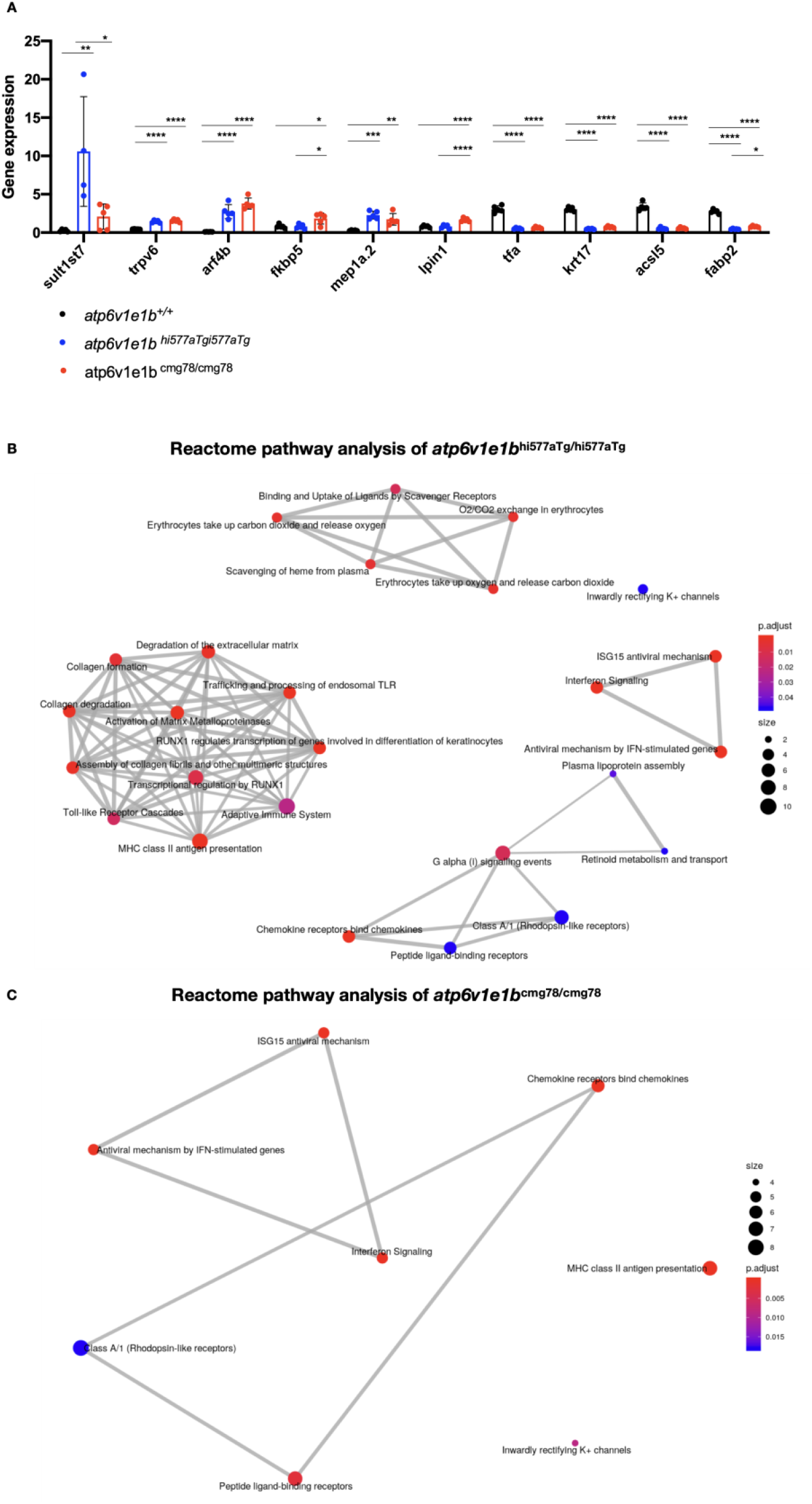
Reactome pathway analysis confirms clustering of oxygen exchange, ion delivery and inflammatory signaling. (A) RT-qPCR analysis of 10 from the top150 DEGs validated the results obtained in the unbiased transcriptomic analysis. (B-C) Cluster plot showing the top enriched reactome pathways of the DEG from whole-body samples of *atp6v1e1b*-deficient and WT zebrafish at 3 dpf.

**Supplemental Figure 4:**
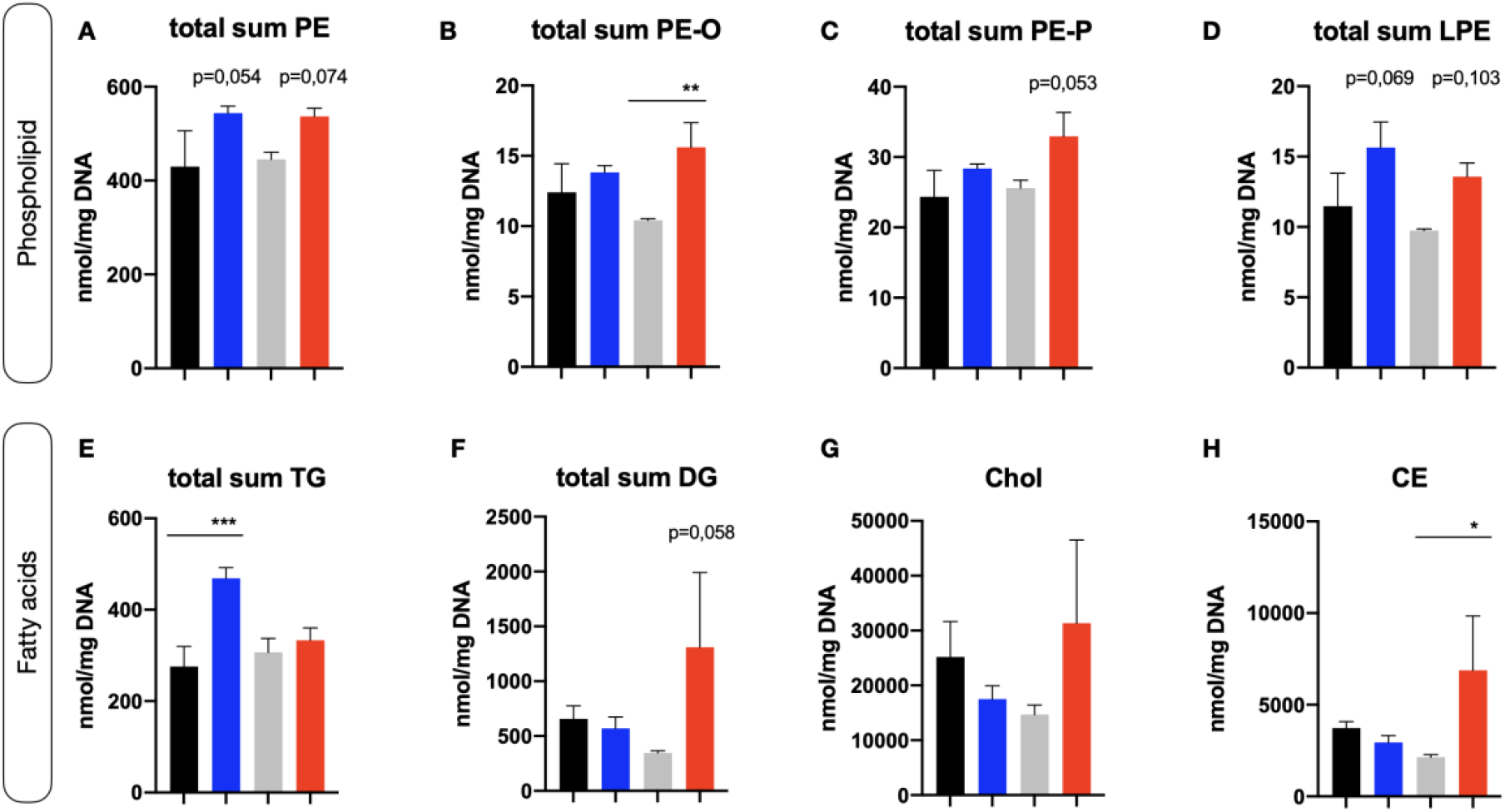
Minor changes in phospholipid and fatty acid levels upon *atp6v1e1b* depletion in zebrafish larvae. (A-H) HILIC LC MS/MS lipidomic analysis demonstrates minor changes in the total levels of phospholipids, and fatty acids in 3 dpf *atp6v1e1b*-deficient zebrafish. Data are presented as mean ± SD from 3 biological replicates. TG: triacylglycerides; DG: diacylglycerides; Chol: cholesterol; CE: cholesterol esters; PE: phosphatidylethanolamine; PE-O: 1-alkyl,2-acylphosphatidylethanolamines; PE-P: 1-alkenyl,2-acylphosphatidylethanolamines; LPE: lysophosphatidylethanolamine.

**Supplemental Figure 5:**
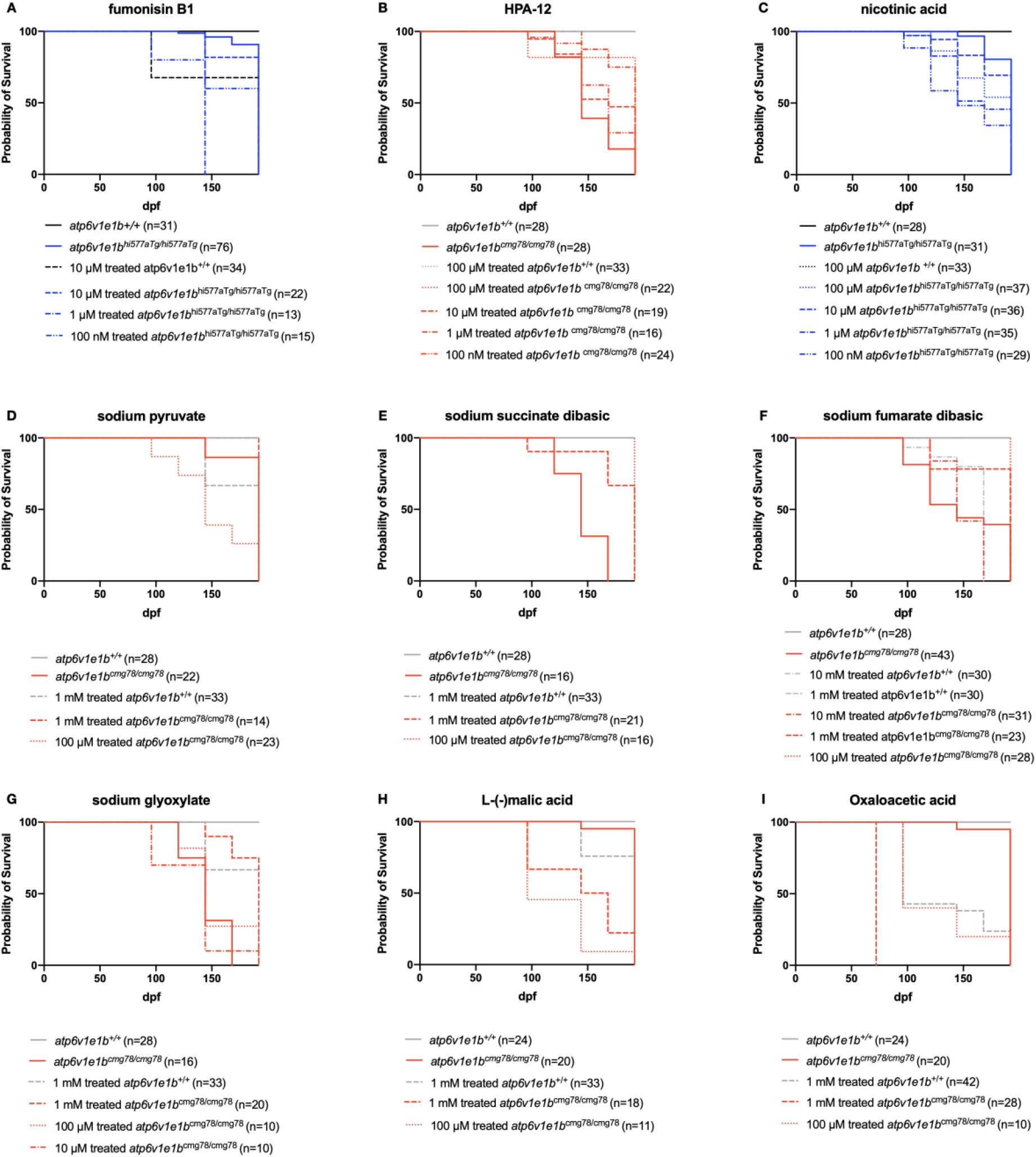
Modulating the ceramide synthesis pathway or the TCA cycle does not ameliorate the survival of *atp6v1e1b*-deficient zebrafish. **A-I**) Survival curves are shown for *atp6v1e1b*-deficient zebrafish and controls until 8 dpf. Compounds were administered at 1 dpf, after chorion removal. A-C) Administration of compounds interfering with the ceramide biosynthesis and cholesterol esterification. D-I) Administration of compounds influencing the TCA cycle. Kaplan-Meier curves for survival of *atp6v1e1b*-deficient and treated *atp6v1e1b*-deficient zebrafish.

**Supplemental Table 1:** List of DEGs with unknown gene names.

| Ensemble | ZFIN | Chromosome | Predicted gene association |
| --- | --- | --- | --- |
| CU984600.2 | si:dkeyp-118h9.7 | Chromosome 9:<br>563,547-582,037<br>forward strand | GTPBP2 (HGNC<br>Symbol) |
| si:ch1073-340i21.3 | si:ch1073-340i21.3 | Chromosome 15:<br>46,329,149-46,336,399 forward<br>strand. | *fb16f09* |
| si:dkey-35h6.1 | si:dkey-35h6.1 | Chromosome 14:<br>13,036,560-13,048,355 reverse<br>strand. | NA |

**Supplemental Table 2:**
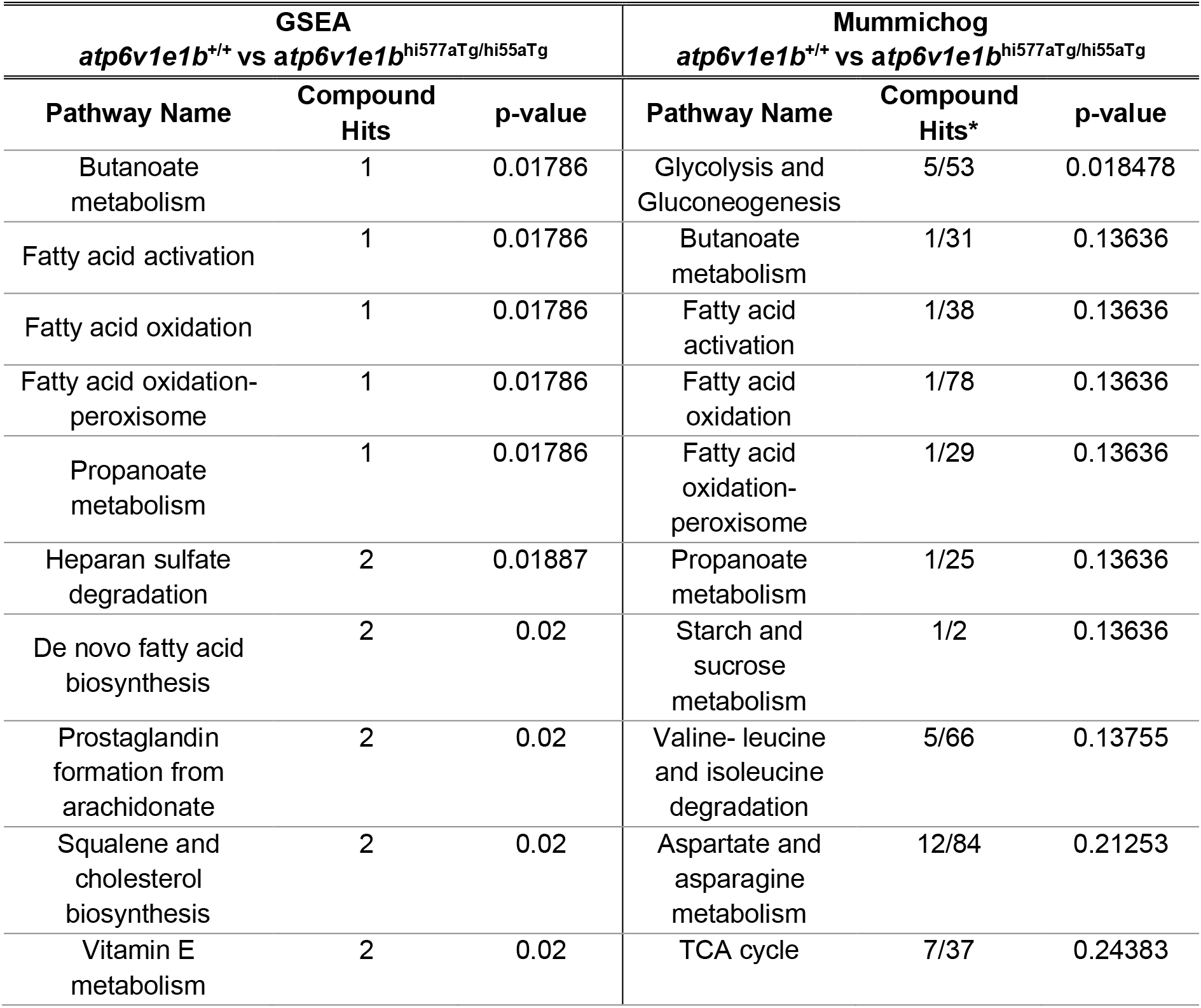
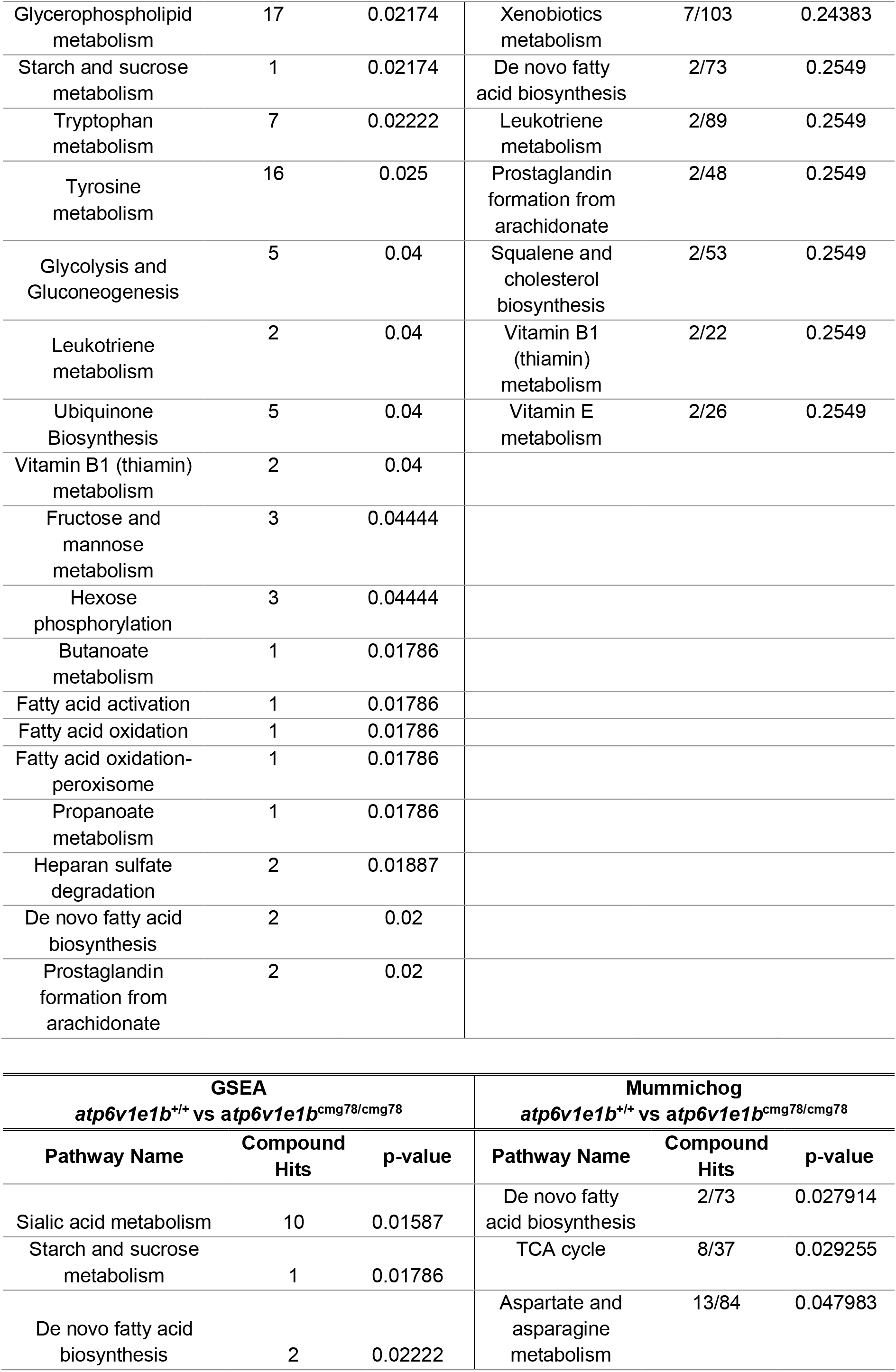
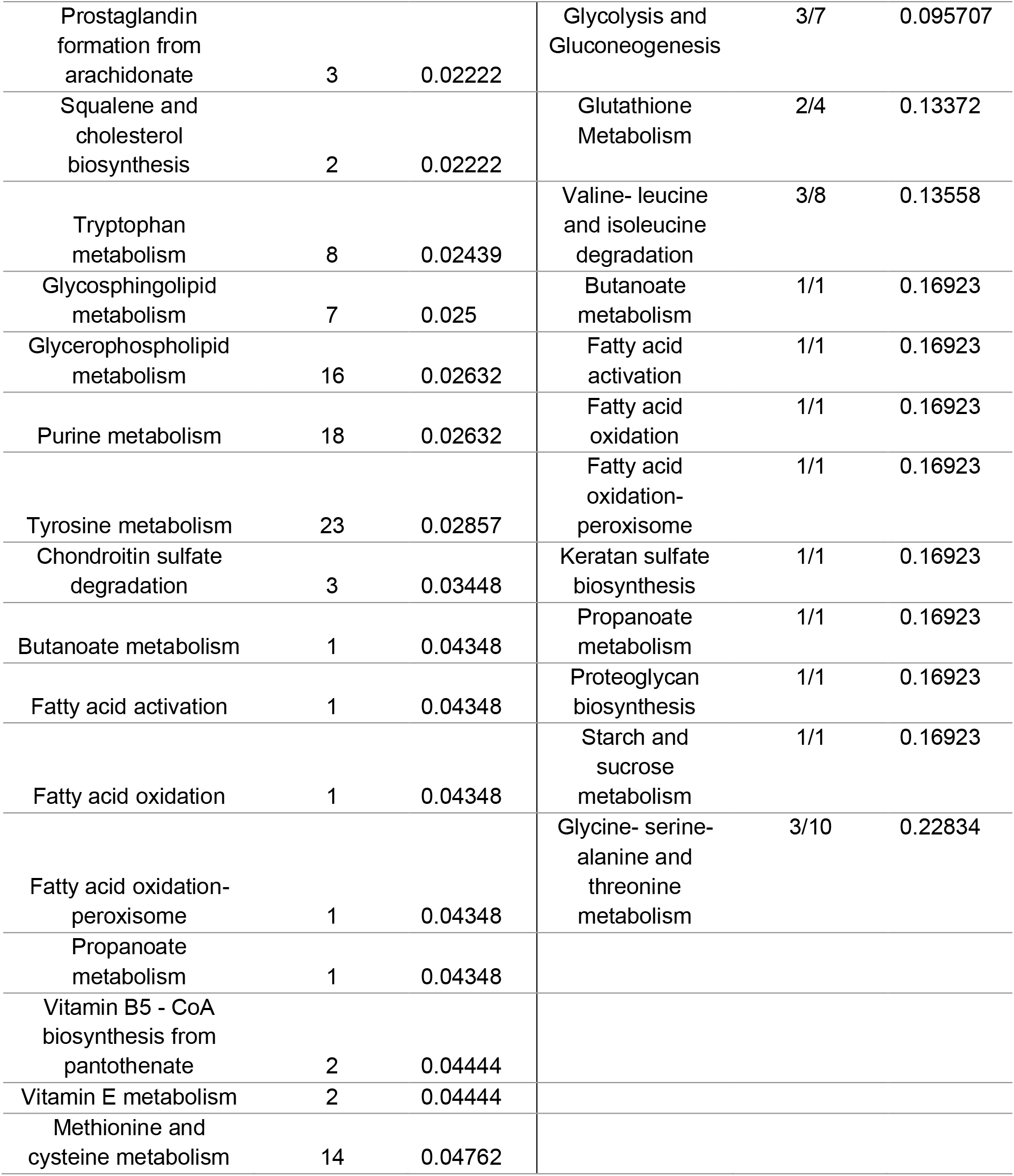
Pathway activity profile of metabolites from *atp6v1e1b*-deficient zebrafish. (Left) GSEA algorithm which considers the overall ranks of features without using a significance cut-off. (Right) Mummichog algorithm which implements an over-representation analysis method that is not able to detect subtle changes. *The mummichog compound hits represent the number of significant compounds divided by the total number of compounds per pathway.

**Supplemental Table 3:**
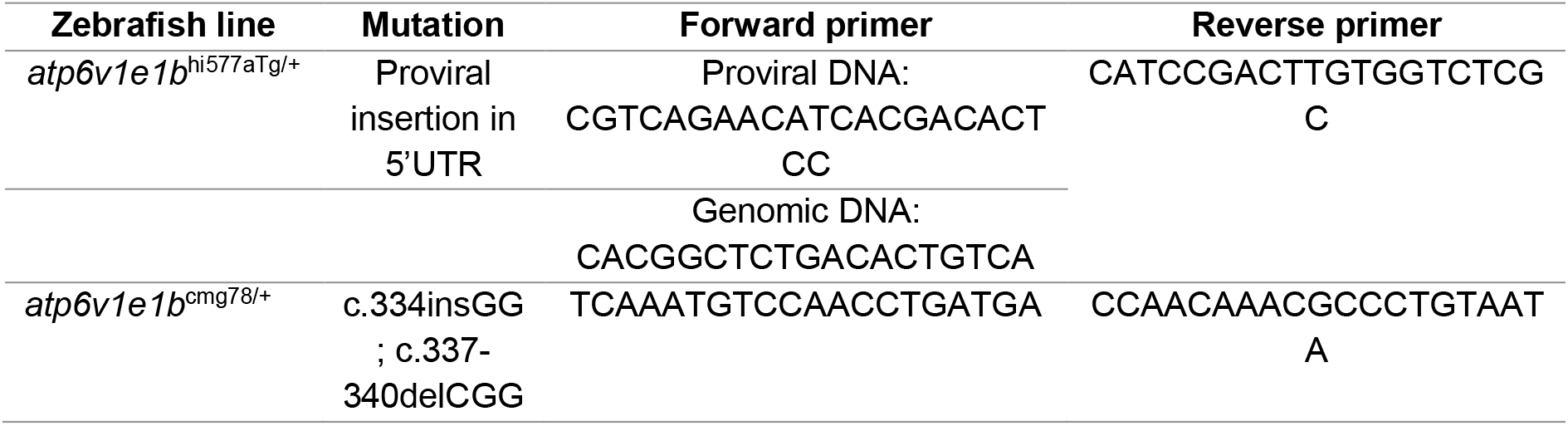
Zebrafish mutation and primers used for genotyping.

**Supplemental Table 4:** Compound screen in zebrafish.

| Compound | Supplier | Cat. No. |
| --- | --- | --- |
| Sodium fumarate dibasic | Sigma-Aldrich | F1506 |
| Sodium pyruvate | Sigma-Aldrich | P2256 |
| Oxaloacetic acid | Sigma-Aldrich | O4126 |
| L-(-)-Malic acid | Sigma-Aldrich | M1000 |
| Sodium succinate dibasic hexahydrate | Sigma-Aldrich | F1506 |
| Sodium glyoxylate monohydrate | Sigma-Aldrich | G4502 |
| Fumonisin B1 | R&D Systems Europe | 3103/1 |
| HPA-12 | TCI Europe NV | H1553 |
| Nicotinic acid | Sigma-Aldrich | N0761 |
| Ammonium iron(III) citrate | Sigma-Aldrich | F5879 |

**Supplemental Table 5:**
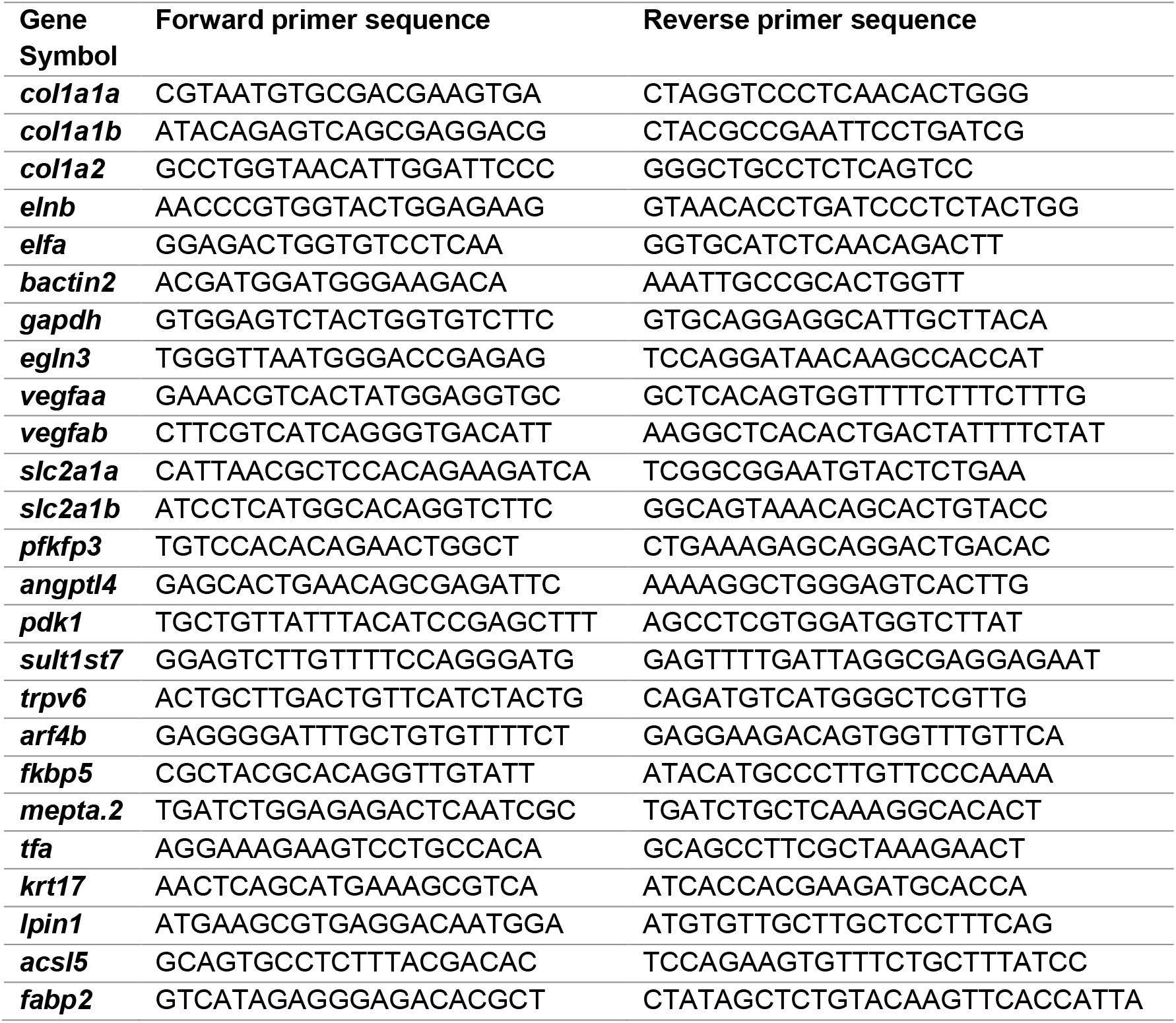
Primer sequences for qPCR analysis.

## Notes

**Funding** B.C. is a senior clinical investigator of the Research Foundation Flanders. This work was supported by a starting grant of the Special Research Fund, Flanders of Ghent University (grant 01N04516C to B.C.), by a Methusalem Grant (BOFMET2015000401) from Ghent University, by a junior fundamental research project grant of the Fund for Scientific Research (G035620N) and by the European Union’s Horizon 2020 research and innovation program, under the Marie Skłodowska-Curie grant agreement No. 794365 to P.S.

https://www.ebi.ac.uk/arrayexpress/experiments/E-MTAB-8824

### Competing Interest Statement

The authors have declared no competing interest.

## References

1. Collins MP, Forgac M. Regulation and function of V-ATPases in physiology and disease. Biochimica et biophysica acta Biomembranes. 2020:183341. Epub 2020/05/19. doi: 10.1016/j.bbamem.2020.183341. PubMed PMID: 32422136.

2. Korvatska O, Strand NS, Berndt JD, Strovas T, Chen DH, Leverenz JB, et al. Altered splicing of ATP6AP2 causes X-linked parkinsonism with spasticity (XPDS). Human molecular genetics. 2013;22(16):3259–68. Epub 2013/04/19. doi: 10.1093/hmg/ddt180. PubMed PMID: 23595882; PubMed Central PMCID: PMCPMC3723311.

3. Hedera P, Rainier S, Zhao XP, Schalling M, Lindblad K, Yuan QP, et al. Spastic paraplegia, ataxia, mental retardation (SPAR): a novel genetic disorder. Neurology. 2002;58(3):411–6. Epub 2002/02/13. doi: 10.1212/wnl.58.3.411. PubMed PMID: 11839840.

4. Ramser J, Abidi FE, Burckle CA, Lenski C, Toriello H, Wen G, et al. A unique exonic splice enhancer mutation in a family with X-linked mental retardation and epilepsy points to a novel role of the renin receptor. Human molecular genetics. 2005;14(8):1019–27. Epub 2005/03/05. doi: 10.1093/hmg/ddi094. PubMed PMID: 15746149.

5. Fassio A, Esposito A, Kato M, Saitsu H, Mei D, Marini C, et al. De novo mutations of the ATP6V1A gene cause developmental encephalopathy with epilepsy. Brain : a journal of neurology. 2018;141(6):1703–18. Epub 2018/04/19. doi: 10.1093/brain/awy092. PubMed PMID: 29668857; PubMed Central PMCID: PMCPMC5972584.

6. Rujano MA, Cannata Serio M, Panasyuk G, Peanne R, Reunert J, Rymen D, et al. Mutations in the X-linked ATP6AP2 cause a glycosylation disorder with autophagic defects. The Journal of experimental medicine. 2017;214(12):3707–29. Epub 2017/11/12. doi: 10.1084/jem.20170453. PubMed PMID: 29127204; PubMed Central PMCID: PMCPMC5716037.

7. Jansen EJ, Timal S, Ryan M, Ashikov A, van Scherpenzeel M, Graham LA, et al. ATP6AP1 deficiency causes an immunodeficiency with hepatopathy, cognitive impairment and abnormal protein glycosylation. Nature communications. 2016;7:11600. Epub 2016/05/28. doi: 10.1038/ncomms11600. PubMed PMID: 27231034; PubMed Central PMCID: PMCPMC4894975.

8. Stover EH, Borthwick KJ, Bavalia C, Eady N, Fritz DM, Rungroj N, et al. Novel ATP6V1B1 and ATP6V0A4 mutations in autosomal recessive distal renal tubular acidosis with new evidence for hearing loss. Journal of medical genetics. 2002;39(11):796–803. Epub 2002/11/05. doi: 10.1136/jmg.39.11.796. PubMed PMID: 12414817; PubMed Central PMCID: PMCPMC1735017.

9. Karet FE, Finberg KE, Nelson RD, Nayir A, Mocan H, Sanjad SA, et al. Mutations in the gene encoding B1 subunit of H+-ATPase cause renal tubular acidosis with sensorineural deafness. Nature genetics. 1999;21(1):84–90. Epub 1999/01/23. doi: 10.1038/5022. PubMed PMID: 9916796.

10. Van Damme T, Gardeitchik T, Mohamed M, Guerrero-Castillo S, Freisinger P, Guillemyn B, et al. Mutations in ATP6V1E1 or ATP6V1A Cause Autosomal-Recessive Cutis Laxa. American journal of human genetics. 2017;100(2):216–27. Epub 2017/01/10. doi: 10.1016/j.ajhg.2016.12.010. PubMed PMID: 28065471; PubMed Central PMCID: PMCPMC5294668.

11. Beyens A, Moreno-Artero E, Bodemer C, Cox H, Gezdirici A, Yilmaz Gulec E, et al. ATP6V0A2-related cutis laxa in 10 novel patients: Focus on clinical variability and expansion of the phenotype. Experimental dermatology. 2019;28(10):1142–5. Epub 2018/06/29. doi: 10.1111/exd.13723. PubMed PMID: 29952037.

12. Poorkaj P, Raskind WH, Leverenz JB, Matsushita M, Zabetian CP, Samii A, et al. A novel X-linked four-repeat tauopathy with Parkinsonism and spasticity. Movement disorders : official journal of the Movement Disorder Society. 2010;25(10):1409–17. Epub 2010/07/16. doi: 10.1002/mds.23085. PubMed PMID: 20629132; PubMed Central PMCID: PMCPMC3123999.

13. Guillard M, Dimopoulou A, Fischer B, Morava E, Lefeber DJ, Kornak U, et al. Vacuolar H+-ATPase meets glycosylation in patients with cutis laxa. Biochimica et biophysica acta. 2009;1792(9):903–14. Epub 2009/01/28. doi: 10.1016/j.bbadis.2008.12.009. PubMed PMID: 19171192.

14. Kornak U, Reynders E, Dimopoulou A, van Reeuwijk J, Fischer B, Rajab A, et al. Impaired glycosylation and cutis laxa caused by mutations in the vesicular H+-ATPase subunit ATP6V0A2. Nature genetics. 2008;40(1):32–4. Epub 2007/12/25. doi: 10.1038/ng.2007.45. PubMed PMID: 18157129.

15. Alazami AM, Al-Qattan SM, Faqeih E, Alhashem A, Alshammari M, Alzahrani F, et al. Expanding the clinical and genetic heterogeneity of hereditary disorders of connective tissue. Human genetics. 2016;135(5):525–40. Epub 2016/03/30. doi: 10.1007/s00439-016-1660-z. PubMed PMID: 27023906.

16. Fischer B, Dimopoulou A, Egerer J, Gardeitchik T, Kidd A, Jost D, et al. Further characterization of ATP6V0A2-related autosomal recessive cutis laxa. Human genetics. 2012;131(11):1761–73. Epub 2012/07/10. doi: 10.1007/s00439-012-1197-8. PubMed PMID: 22773132.

17. Cotter K, Stransky L, McGuire C, Forgac M. Recent Insights into the Structure, Regulation, and Function of the V-ATPases. Trends in biochemical sciences. 2015;40(10):611–22. Epub 2015/09/28. doi: 10.1016/j.tibs.2015.08.005. PubMed PMID: 26410601; PubMed Central PMCID: PMCPMC4589219.

18. Stransky L, Cotter K, Forgac M. The Function of V-ATPases in Cancer. Physiological reviews. 2016;96(3):1071–91. Epub 2016/06/24. doi: 10.1152/physrev.00035.2015. PubMed PMID: 27335445; PubMed Central PMCID: PMCPMC4982037.

19. Sennoune SR, Martinez-Zaguilan R. Vacuolar H(+)-ATPase signaling pathway in cancer. Current protein & peptide science. 2012;13(2):152–63. Epub 2011/11/03. doi: 10.2174/138920312800493197. PubMed PMID: 22044157.

20. Kopan R, Ilagan MX. The canonical Notch signaling pathway: unfolding the activation mechanism. Cell. 2009;137(2):216–33. Epub 2009/04/22. doi: 10.1016/j.cell.2009.03.045. PubMed PMID: 19379690; PubMed Central PMCID: PMCPMC2827930.

21. Whitton B, Okamoto H, Packham G, Crabb SJ. Vacuolar ATPase as a potential therapeutic target and mediator of treatment resistance in cancer. Cancer medicine. 2018;7(8):3800–11. Epub 2018/06/22. doi: 10.1002/cam4.1594. PubMed PMID: 29926527; PubMed Central PMCID: PMCPMC6089187.

22. Pakkiriswami S, Couto A, Nagarajan U, Georgiou M. Glycosylated Notch and Cancer. Frontiers in oncology. 2016;6:37. Epub 2016/03/01. doi: 10.3389/fonc.2016.00037. PubMed PMID: 26925390; PubMed Central PMCID: PMCPMC4757683.

23. Pamarthy S, Kulshrestha A, Katara GK, Beaman KD. The curious case of vacuolar ATPase: regulation of signaling pathways. Molecular cancer. 2018;17(1):41. Epub 2018/02/17. doi: 10.1186/s12943-018-0811-3. PubMed PMID: 29448933; PubMed Central PMCID: PMCPMC5815226.

24. Reya T, Clevers H. Wnt signalling in stem cells and cancer. Nature. 2005;434(7035):843–50. Epub 2005/04/15. doi: 10.1038/nature03319. PubMed PMID: 15829953.

25. He TC, Sparks AB, Rago C, Hermeking H, Zawel L, da Costa LT, et al. Identification of c-MYC as a target of the APC pathway. Science (New York, NY). 1998;281(5382):1509–12. Epub 1998/09/04. doi: 10.1126/science.281.5382.1509. PubMed PMID: 9727977.

26. Shtutman M, Zhurinsky J, Simcha I, Albanese C, D’Amico M, Pestell R, et al. The cyclin D1 gene is a target of the beta-catenin/LEF-1 pathway. Proceedings of the National Academy of Sciences of the United States of America. 1999;96(10):5522–7. Epub 1999/05/13. doi: 10.1073/pnas.96.10.5522. PubMed PMID: 10318916; PubMed Central PMCID: PMCPMC21892.

27. Yamamoto H, Komekado H, Kikuchi A. Caveolin is necessary for Wnt-3a-dependent internalization of LRP6 and accumulation of beta-catenin. Developmental cell. 2006;11(2):213–23. Epub 2006/08/08. doi: 10.1016/j.devcel.2006.07.003. PubMed PMID: 16890161.

28. Jho EH, Zhang T, Domon C, Joo CK, Freund JN, Costantini F. Wnt/beta-catenin/Tcf signaling induces the transcription of Axin2, a negative regulator of the signaling pathway. Molecular and cellular biology. 2002;22(4):1172–83. Epub 2002/01/26. doi: 10.1128/mcb.22.4.1172-1183.2002. PubMed PMID: 11809808; PubMed Central PMCID: PMCPMC134648.

29. Seto ES, Bellen HJ. Internalization is required for proper Wingless signaling in Drosophila melanogaster. The Journal of cell biology. 2006;173(1):95–106. Epub 2006/04/12. doi: 10.1083/jcb.200510123. PubMed PMID: 16606693; PubMed Central PMCID: PMCPMC2063794.

30. Yambire KF, Rostosky C, Watanabe T, Pacheu-Grau D, Torres-Odio S, Sanchez-Guerrero A, et al. Impaired lysosomal acidification triggers iron deficiency and inflammation in vivo. eLife. 2019;8. Epub 2019/12/04. doi: 10.7554/eLife.51031. PubMed PMID: 31793879; PubMed Central PMCID: PMCPMC6917501.

31. Miles AL, Burr SP, Grice GL, Nathan JA. The vacuolar-ATPase complex and assembly factors, TMEM199 and CCDC115, control HIF1alpha prolyl hydroxylation by regulating cellular iron levels. eLife. 2017;6. Epub 2017/03/16. doi: 10.7554/eLife.22693. PubMed PMID: 28296633; PubMed Central PMCID: PMCPMC5391204.

32. Weber RA, Yen FS, Nicholson SPV, Alwaseem H, Bayraktar EC, Alam M, et al. Maintaining Iron Homeostasis Is the Key Role of Lysosomal Acidity for Cell Proliferation. Molecular cell. 2020;77(3):645–55 e7. Epub 2020/01/28. doi: 10.1016/j.molcel.2020.01.003. PubMed PMID: 31983508; PubMed Central PMCID: PMCPMC7176020.

33. Sun-Wada G, Murata Y, Yamamoto A, Kanazawa H, Wada Y, Futai M. Acidic endomembrane organelles are required for mouse postimplantation development. Developmental biology. 2000;228(2):315–25. Epub 2000/12/09. doi: 10.1006/dbio.2000.9963. PubMed PMID: 11112332.

34. Gaiano N, Amsterdam A, Kawakami K, Allende M, Becker T, Hopkins N. Insertional mutagenesis and rapid cloning of essential genes in zebrafish. Nature. 1996;383(6603):829–32. Epub 1996/10/31. doi: 10.1038/383829a0. PubMed PMID: 8893009.

35. Nuckels RJ, Ng A, Darland T, Gross JM. The vacuolar-ATPase complex regulates retinoblast proliferation and survival, photoreceptor morphogenesis, and pigmentation in the zebrafish eye. Investigative ophthalmology & visual science. 2009;50(2):893–905. Epub 2008/10/07. doi: 10.1167/iovs.08-2743. PubMed PMID: 18836173.

36. Yamakawa N, Vanbeselaere J, Chang LY, Yu SY, Ducrocq L, Harduin-Lepers A, et al. Systems glycomics of adult zebrafish identifies organ-specific sialylation and glycosylation patterns. Nature communications. 2018;9(1):4647. Epub 2018/11/09. doi: 10.1038/s41467-018-06950-3. PubMed PMID: 30405127; PubMed Central PMCID: PMCPMC6220181.

37. Guerardel Y, Chang LY, Maes E, Huang CJ, Khoo KH. Glycomic survey mapping of zebrafish identifies unique sialylation pattern. Glycobiology. 2006;16(3):244–57. Epub 2005/12/03. doi: 10.1093/glycob/cwj062. PubMed PMID: 16321922.

38. Hanzawa K, Suzuki N, Natsuka S. Structures and developmental alterations of N-glycans of zebrafish embryos. Glycobiology. 2017;27(3):228–45. Epub 2016/12/10. doi: 10.1093/glycob/cww124. PubMed PMID: 27932382.

39. Ofer S, Fibach E, Kessel M, Bauminger ER, Cohen SG, Eikelboom J, et al. Iron incorporation into ferritin and hemoglobin during differentiation of murine erythroleukemia cells. Blood. 1981;58(2):255–62. Epub 1981/08/01. PubMed PMID: 6941820.

40. Mookerjee SA, Gerencser AA, Nicholls DG, Brand MD. Quantifying intracellular rates of glycolytic and oxidative ATP production and consumption using extracellular flux measurements. The Journal of biological chemistry. 2017;292(17):7189–207. Epub 2017/03/09. doi: 10.1074/jbc.M116.774471. PubMed PMID: 28270511; PubMed Central PMCID: PMCPMC5409486.

41. Symersky J, Osowski D, Walters DE, Mueller DM. Oligomycin frames a common drug-binding site in the ATP synthase. Proceedings of the National Academy of Sciences of the United States of America. 2012;109(35):13961–5. Epub 2012/08/08. doi: 10.1073/pnas.1207912109. PubMed PMID: 22869738; PubMed Central PMCID: PMCPMC3435195.

42. Udono M, Fujii K, Harada G, Tsuzuki Y, Kadooka K, Zhang P, et al. Impaired ATP6V0A2 expression contributes to Golgi dispersion and glycosylation changes in senescent cells. Scientific reports. 2015;5:17342. Epub 2015/11/28. doi: 10.1038/srep17342. PubMed PMID: 26611489; PubMed Central PMCID: PMCPMC4661525.

43. Page MGP. The Role of Iron and Siderophores in Infection, and the Development of Siderophore Antibiotics. Clinical infectious diseases : an official publication of the Infectious Diseases Society of America. 2019;69(Suppl 7):S529–S37. Epub 2019/11/15. doi: 10.1093/cid/ciz825. PubMed PMID: 31724044; PubMed Central PMCID: PMCPMC6853763.

44. Napier I, Ponka P, Richardson DR. Iron trafficking in the mitochondrion: novel pathways revealed by disease. Blood. 2005;105(5):1867–74. Epub 2004/11/06. doi: 10.1182/blood-2004-10-3856. PubMed PMID: 15528311.

45. Fraenkel PG, Gibert Y, Holzheimer JL, Lattanzi VJ, Burnett SF, Dooley KA, et al. Transferrin-a modulates hepcidin expression in zebrafish embryos. Blood. 2009;113(12):2843–50. Epub 2008/12/03. doi: 10.1182/blood-2008-06-165340. PubMed PMID: 19047682; PubMed Central PMCID: PMCPMC2661867.

46. Devireddy LR, Hart DO, Goetz DH, Green MR. A mammalian siderophore synthesized by an enzyme with a bacterial homolog involved in enterobactin production. Cell. 2010;141(6):1006–17. Epub 2010/06/17. doi: 10.1016/j.cell.2010.04.040. PubMed PMID: 20550936; PubMed Central PMCID: PMCPMC2910436.

47. Qi B, Han M. Microbial Siderophore Enterobactin Promotes Mitochondrial Iron Uptake and Development of the Host via Interaction with ATP Synthase. Cell. 2018;175(2):571–82 e11. Epub 2018/08/28. doi: 10.1016/j.cell.2018.07.032. PubMed PMID: 30146159.

48. Chen CT, Hsu SH, Wei YH. Upregulation of mitochondrial function and antioxidant defense in the differentiation of stem cells. Biochimica et biophysica acta. 2010;1800(3):257–63. Epub 2009/09/15. doi: 10.1016/j.bbagen.2009.09.001. PubMed PMID: 19747960.

49. Parkinson-Lawrence EJ, Shandala T, Prodoehl M, Plew R, Borlace GN, Brooks DA. Lysosomal storage disease: revealing lysosomal function and physiology. Physiology (Bethesda, Md). 2010;25(2):102–15. Epub 2010/05/01. doi: 10.1152/physiol.00041.2009. PubMed PMID: 20430954.

50. Platt FM, Boland B, van der Spoel AC. The cell biology of disease: lysosomal storage disorders: the cellular impact of lysosomal dysfunction. The Journal of cell biology. 2012;199(5):723–34. Epub 2012/11/28. doi: 10.1083/jcb.201208152. PubMed PMID: 23185029; PubMed Central PMCID: PMCPMC3514785.

51. Zlamy M, Hofstatter J, Albrecht U, Baumgartner S, Haberlandt E, Scholl-Burgi S, et al. The value of axillary skin electron microscopic analysis in the diagnosis of lysosomal storage disorders. Modern pathology : an official journal of the United States and Canadian Academy of Pathology, Inc. 2019;32(6):755–63. Epub 2019/02/07. doi: 10.1038/s41379-019-0201-4. PubMed PMID: 30723298.

52. Lafourcade C, Sobo K, Kieffer-Jaquinod S, Garin J, van der Goot FG. Regulation of the V-ATPase along the endocytic pathway occurs through reversible subunit association and membrane localization. PloS one. 2008;3(7):e2758. Epub 2008/07/24. doi: 10.1371/journal.pone.0002758. PubMed PMID: 18648502; PubMed Central PMCID: PMCPMC2447177.

53. Sobo K, Chevallier J, Parton RG, Gruenberg J, van der Goot FG. Diversity of raft-like domains in late endosomes. PloS one. 2007;2(4):e391. Epub 2007/04/27. doi: 10.1371/journal.pone.0000391. PubMed PMID: 17460758; PubMed Central PMCID: PMCPMC1851096.

54. Huotari J, Helenius A. Endosome maturation. The EMBO journal. 2011;30(17):3481–500. Epub 2011/09/01. doi: 10.1038/emboj.2011.286. PubMed PMID: 21878991; PubMed Central PMCID: PMCPMC3181477.

55. Hannun YA, Obeid LM. Sphingolipids and their metabolism in physiology and disease. Nature reviews Molecular cell biology. 2018;19(3):175–91. Epub 2017/11/23. doi: 10.1038/nrm.2017.107. PubMed PMID: 29165427; PubMed Central PMCID: PMCPMC5902181.

56. Tabassum R, Ramo JT, Ripatti P, Koskela JT, Kurki M, Karjalainen J, et al. Genetic architecture of human plasma lipidome and its link to cardiovascular disease. Nature communications. 2019;10(1):4329. Epub 2019/09/26. doi: 10.1038/s41467-019-11954-8. PubMed PMID: 31551469; PubMed Central PMCID: PMCPMC6760179.

57. Khavandgar Z, Murshed M. Sphingolipid metabolism and its role in the skeletal tissues. Cellular and molecular life sciences : CMLS. 2015;72(5):959–69. Epub 2014/11/27. doi: 10.1007/s00018-014-1778-x. PubMed PMID: 25424644.

58. Campanella M, Parker N, Tan CH, Hall AM, Duchen MR. IF(1): setting the pace of the F(1)F(o)-ATP synthase. Trends in biochemical sciences. 2009;34(7):343–50. Epub 2009/06/30. doi: 10.1016/j.tibs.2009.03.006. PubMed PMID: 19559621.

59. Campanella M, Casswell E, Chong S, Farah Z, Wieckowski MR, Abramov AY, et al. Regulation of mitochondrial structure and function by the F1Fo-ATPase inhibitor protein, IF1. Cell metabolism. 2008;8(1):13–25. Epub 2008/07/02. doi: 10.1016/j.cmet.2008.06.001. PubMed PMID: 18590689.

60. Baker N, Hamilton G, Wilkes JM, Hutchinson S, Barrett MP, Horn D. Vacuolar ATPase depletion affects mitochondrial ATPase function, kinetoplast dependency, and drug sensitivity in trypanosomes. Proceedings of the National Academy of Sciences of the United States of America. 2015;112(29):9112–7. Epub 2015/07/08. doi: 10.1073/pnas.1505411112. PubMed PMID: 26150481; PubMed Central PMCID: PMCPMC4517229.

61. Hinton A, Sennoune SR, Bond S, Fang M, Reuveni M, Sahagian GG, et al. Function of a subunit isoforms of the V-ATPase in pH homeostasis and in vitro invasion of MDA-MB231 human breast cancer cells. The Journal of biological chemistry. 2009;284(24):16400–8. Epub 2009/04/16. doi: 10.1074/jbc.M901201200. PubMed PMID: 19366680; PubMed Central PMCID: PMCPMC2713521.

62. Capecci J, Forgac M. The function of vacuolar ATPase (V-ATPase) a subunit isoforms in invasiveness of MCF10a and MCF10CA1a human breast cancer cells. The Journal of biological chemistry. 2013;288(45):32731–41. Epub 2013/09/28. doi: 10.1074/jbc.M113.503771. PubMed PMID: 24072707; PubMed Central PMCID: PMCPMC3820907.

63. McGuire CM, Collins MP, Sun-Wada G, Wada Y, Forgac M. Isoform-specific gene disruptions reveal a role for the V-ATPase subunit a4 isoform in the invasiveness of 4T1-12B breast cancer cells. The Journal of biological chemistry. 2019;294(29):11248–58. Epub 2019/06/07. doi: 10.1074/jbc.RA119.007713. PubMed PMID: 31167791; PubMed Central PMCID: PMCPMC6643023.

64. De Milito A, Canese R, Marino ML, Borghi M, Iero M, Villa A, et al. pH-dependent antitumor activity of proton pump inhibitors against human melanoma is mediated by inhibition of tumor acidity. International journal of cancer. 2010;127(1):207–19. Epub 2009/10/31. doi: 10.1002/ijc.25009. PubMed PMID: 19876915.

65. Neri D, Supuran CT. Interfering with pH regulation in tumours as a therapeutic strategy. Nature reviews Drug discovery. 2011;10(10):767–77. Epub 2011/09/17. doi: 10.1038/nrd3554. PubMed PMID: 21921921.

66. Gottlieb RA, Giesing HA, Zhu JY, Engler RL, Babior BM. Cell acidification in apoptosis: granulocyte colony-stimulating factor delays programmed cell death in neutrophils by up-regulating the vacuolar H(+)-ATPase. Proceedings of the National Academy of Sciences of the United States of America. 1995;92(13):5965–8. Epub 1995/06/20. doi: 10.1073/pnas.92.13.5965. PubMed PMID: 7541139; PubMed Central PMCID: PMCPMC41622.

67. Kallifatidis G, Hoepfner D, Jaeg T, Guzman EA, Wright AE. The marine natural product manzamine A targets vacuolar ATPases and inhibits autophagy in pancreatic cancer cells. Marine drugs. 2013;11(9):3500–16. Epub 2013/09/21. doi: 10.3390/md11093500. PubMed PMID: 24048269; PubMed Central PMCID: PMCPMC3806460.

68. Karwatowska-Prokopczuk E, Nordberg JA, Li HL, Engler RL, Gottlieb RA. Effect of vacuolar proton ATPase on pHi, Ca2+, and apoptosis in neonatal cardiomyocytes during metabolic inhibition/recovery. Circulation research. 1998;82(11):1139–44. Epub 1998/06/20. doi: 10.1161/01.res.82.11.1139. PubMed PMID: 9633914.

69. Amsterdam A, Nissen RM, Sun Z, Swindell EC, Farrington S, Hopkins N. Identification of 315 genes essential for early zebrafish development. Proceedings of the National Academy of Sciences of the United States of America. 2004;101(35):12792–7. Epub 2004/07/17. doi: 10.1073/pnas.0403929101. PubMed PMID: 15256591; PubMed Central PMCID: PMCPMC516474.

70. Boel A, Steyaert W, De Rocker N, Menten B, Callewaert B, De Paepe A, et al. BATCH-GE: Batch analysis of Next-Generation Sequencing data for genome editing assessment. Scientific reports. 2016;6:30330. Epub 2016/07/28. doi: 10.1038/srep30330. PubMed PMID: 27461955; PubMed Central PMCID: PMCPMC4962088.

71. Lawrence C, Sanders GE, Varga ZM, Baumann DP, Freeman A, Baur B, et al. Regulatory compliance and the zebrafish. Zebrafish. 2009;6(4):453–6. Epub 2009/11/18. doi: 10.1089/zeb.2009.0595. PubMed PMID: 19916799.

72. Westerfield M, Doerry E, Kirkpatrick AE, Douglas SA. Zebrafish informatics and the ZFIN database. Methods in cell biology. 1999;60:339–55. Epub 1999/01/19. doi: 10.1016/s0091-679x(08)61909-3. PubMed PMID: 9891346.

73. Schindelin J, Arganda-Carreras I, Frise E, Kaynig V, Longair M, Pietzsch T, et al. Fiji: an open-source platform for biological-image analysis. Nature methods. 2012;9(7):676–82. Epub 2012/06/30. doi: 10.1038/nmeth.2019. PubMed PMID: 22743772; PubMed Central PMCID: PMCPMC3855844.

74. Wang M, Sips P, Khin E, Rotival M, Sun X, Ahmed R, et al. Wars2 is a determinant of angiogenesis. Nature communications. 2016;7:12061. Epub 2016/07/09. doi: 10.1038/ncomms12061. PubMed PMID: 27389904; PubMed Central PMCID: PMCPMC4941120.

75. Shin JT, Pomerantsev EV, Mably JD, MacRae CA. High-resolution cardiovascular function confirms functional orthology of myocardial contractility pathways in zebrafish. Physiological genomics. 2010;42(2):300–9. Epub 2010/04/15. doi: 10.1152/physiolgenomics.00206.2009. PubMed PMID: 20388839; PubMed Central PMCID: PMCPMC3032279.

76. Gistelinck C, Kwon RY, Malfait F, Symoens S, Harris MP, Henke K, et al. Zebrafish type I collagen mutants faithfully recapitulate human type I collagenopathies. Proceedings of the National Academy of Sciences of the United States of America. 2018;115(34):E8037–E46. Epub 2018/08/08. doi: 10.1073/pnas.1722200115. PubMed PMID: 30082390; PubMed Central PMCID: PMCPMC6112716.

77. Isogai S, Horiguchi M, Weinstein BM. The vascular anatomy of the developing zebrafish: an atlas of embryonic and early larval development. Developmental biology. 2001;230(2):278–301. Epub 2001/02/13. doi: 10.1006/dbio.2000.9995. PubMed PMID: 11161578.

78. Huysseune A, Sire JY. Bone and cartilage resorption in relation to tooth development in the anterior part of the mandible in cichlid fish: a light and TEM study. The Anatomical record. 1992;234(1):1–14. Epub 1992/09/01. doi: 10.1002/ar.1092340102. PubMed PMID: 1416089.

79. Gistelinck C, Witten PE, Huysseune A, Symoens S, Malfait F, Larionova D, et al. Loss of Type I Collagen Telopeptide Lysyl Hydroxylation Causes Musculoskeletal Abnormalities in a Zebrafish Model of Bruck Syndrome. Journal of bone and mineral research : the official journal of the American Society for Bone and Mineral Research. 2016;31(11):1930–42. Epub 2016/10/25. doi: 10.1002/jbmr.2977. PubMed PMID: 27541483; PubMed Central PMCID: PMCPMC5364950.

80. Ibhazehiebo K, Gavrilovici C, de la Hoz CL, Ma SC, Rehak R, Kaushik G, et al. A novel metabolism-based phenotypic drug discovery platform in zebrafish uncovers HDACs 1 and 3 as a potential combined anti-seizure drug target. Brain : a journal of neurology. 2018;141(3):744–61. Epub 2018/01/27. doi: 10.1093/brain/awx364. PubMed PMID: 29373639; PubMed Central PMCID: PMCPMC5837409.

81. Stackley KD, Beeson CC, Rahn JJ, Chan SS. Bioenergetic profiling of zebrafish embryonic development. PloS one. 2011;6(9):e25652. Epub 2011/10/08. doi: 10.1371/journal.pone.0025652. PubMed PMID: 21980518; PubMed Central PMCID: PMCPMC3183059.

82. Zhang JL, Laurence Souders C, 2nd, Denslow ND, Martyniuk CJ. Quercetin, a natural product supplement, impairs mitochondrial bioenergetics and locomotor behavior in larval zebrafish (Danio rerio). Toxicology and applied pharmacology. 2017;327:30–8. Epub 2017/04/30. doi: 10.1016/j.taap.2017.04.024. PubMed PMID: 28450151.

83. Delbaere S, Van Damme T, Syx D, Symoens S, Coucke P, Willaert A, et al. Hypomorphic zebrafish models mimic the musculoskeletal phenotype of beta4GalT7-deficient Ehlers-Danlos syndrome. Matrix biology : journal of the International Society for Matrix Biology. 2019. Epub 2019/12/22. doi: 10.1016/j.matbio.2019.12.002. PubMed PMID: 31862401.

84. Di Tommaso P, Chatzou M, Floden EW, Barja PP, Palumbo E, Notredame C. Nextflow enables reproducible computational workflows. Nature biotechnology. 2017;35(4):316–9. Epub 2017/04/12. doi: 10.1038/nbt.3820. PubMed PMID: 28398311.

85. Wang L, Wang S, Li W. RSeQC: quality control of RNA-seq experiments. Bioinformatics (Oxford, England). 2012;28(16):2184–5. Epub 2012/06/30. doi: 10.1093/bioinformatics/bts356. PubMed PMID: 22743226.

86. Ewels P, Magnusson M, Lundin S, Kaller M. MultiQC: summarize analysis results for multiple tools and samples in a single report. Bioinformatics (Oxford, England). 2016;32(19):3047–8. Epub 2016/06/18. doi: 10.1093/bioinformatics/btw354. PubMed PMID: 27312411; PubMed Central PMCID: PMCPMC5039924.

87. Dobin A, Davis CA, Schlesinger F, Drenkow J, Zaleski C, Jha S, et al. STAR: ultrafast universal RNA-seq aligner. Bioinformatics (Oxford, England). 2013;29(1):15–21. Epub 2012/10/30. doi: 10.1093/bioinformatics/bts635. PubMed PMID: 23104886; PubMed Central PMCID: PMCPMC3530905.

88. Liao Y, Smyth GK, Shi W. featureCounts: an efficient general purpose program for assigning sequence reads to genomic features. Bioinformatics (Oxford, England). 2014;30(7):923–30. Epub 2013/11/15. doi: 10.1093/bioinformatics/btt656. PubMed PMID: 24227677.

89. Love MI, Huber W, Anders S. Moderated estimation of fold change and dispersion for RNA-seq data with DESeq2. Genome biology. 2014;15(12):550. Epub 2014/12/18. doi: 10.1186/s13059-014-0550-8. PubMed PMID: 25516281; PubMed Central PMCID: PMCPMC4302049.

90. Luo W, Friedman MS, Shedden K, Hankenson KD, Woolf PJ. GAGE: generally applicable gene set enrichment for pathway analysis. BMC bioinformatics. 2009;10:161. Epub 2009/05/29. doi: 10.1186/1471-2105-10-161. PubMed PMID: 19473525; PubMed Central PMCID: PMCPMC2696452.

91. Xia J, Wishart DS. Using MetaboAnalyst 3.0 for Comprehensive Metabolomics Data Analysis. Current protocols in bioinformatics. 2016;55:1401–091. Epub 2016/09/08. doi: 10.1002/cpbi.11. PubMed PMID: 27603023.

92. Chong J, Wishart DS, Xia J. Using MetaboAnalyst 4.0 for Comprehensive and Integrative Metabolomics Data Analysis. Current protocols in bioinformatics. 2019;68(1):e86. Epub 2019/11/23. doi: 10.1002/cpbi.86. PubMed PMID: 31756036.

